# Wnt activity reveals context-specific genetic effects on gene regulation in neural progenitors

**DOI:** 10.1101/2023.02.07.527357

**Authors:** Nana Matoba, Brandon D Le, Jordan M Valone, Justin M Wolter, Jessica Mory, Dan Liang, Nil Aygün, K Alaine Broadaway, Marielle L Bond, Karen L Mohlke, Mark J Zylka, Michael I Love, Jason L Stein

**Author notes:** These authors contributed equally to this work (Nana Matoba, Brandon D Le, Jordan M Valone).

## Abstract

Gene regulatory effects in bulk-post mortem brain tissues are undetected at many non-coding brain trait-associated loci. We hypothesized that context-specific genetic variant function during stimulation of a developmental signaling pathway would explain additional regulatory mechanisms. We measured chromatin accessibility and gene expression following activation of the canonical Wnt pathway in primary human neural progenitors from 82 donors. TCF/LEF motifs, brain structure-, and neuropsychiatric disorder-associated variants were enriched within Wnt-responsive regulatory elements (REs). Genetically influenced REs were enriched in genomic regions under positive selection along the human lineage. Stimulation of the Wnt pathway increased the detection of genetically influenced REs/genes by 66.2%/52.7%, and led to the identification of 397 REs primed for effects on gene expression. Context-specific molecular quantitative trait loci increased brain-trait colocalizations by up to 70%, suggesting that genetic variant effects during early neurodevelopmental patterning lead to differences in adult brain and behavioral traits.

---

Common genetic variation associated with brain-relevant traits and risk for neuropsychiatric disorders have been identified and replicated, providing a molecular basis for understanding inter-individual variation in brain structure, function, and behavior^1, 2^. However, brain-trait associated loci are mostly found in non-coding regions without clear mechanisms of action. Gene regulatory mechanisms of non-coding loci are inferred using datasets mapping the effects of genetic variation on regulatory element activity, marked by accessible chromatin peaks (chromatin accessibility quantitative trait loci or caQTL), or gene expression (eQTL)^3^. Gene regulatory associated loci (ca/eQTLs) measured in bulk post-mortem tissue have explained mechanisms for a subset of brain-trait associated loci through sharing, or colocalization, of causal variants^4–6^. Yet, many brain-trait associated variants do not have detectable gene regulatory function in bulk post-mortem brain tissue, leading to the question of where the ‘missing regulation’ linking trait-associated variants to gene expression lies^4, 7, 8^.

One potential solution is that variants impact the accessibility of regulatory elements or the expression of target genes only in specific contexts or when activated by certain stimuli (response-QTLs), and therefore are unlikely to be observed in bulk post-mortem tissue^9^. The context specificity of genetic variant function, despite identical genetic sequence (excluding somatic mutations) being present in every cell, may be explained in part through the action of transcription factors only expressed, activated, or translocated to the nucleus within certain cell-types or during stimulation. Context-specific genetic effects on gene expression are consistent with the observation that only a subset of genetic variants affecting unstimulated chromatin accessibility also affect gene expression, suggesting that some regulatory elements are primed to impact gene expression when a stimulus leads to the activation of additional transcription factors^9, 10^. Recent studies characterizing cell-type specific ca/eQTLs highlight the importance of cellular context by revealing novel brain trait colocalizations undetected in bulk tissues^10–13^. We hypothesized that the stimulation of a developmental signaling pathway in a homogeneous neural cell type would reveal previously undetected functions of genetic variation and explain some of the ‘missing regulation’ for brain-trait associated loci.

We evaluated context-specific effects of genetic variation in a population of primary human neural progenitor cells (hNPCs), a developmental cell type with regulatory elements enriched for genetic association signals for multiple-brain related traits and neuropsychiatric disorders^10, 11, 14^. We measured chromatin accessibility and gene expression in hNPCs following stimulation of the canonical Wnt pathway, which is known to impact neural progenitor proliferation, cortical patterning, and is associated with the inter-individual differences in complex brain traits^2, 15–23^. Wnt stimulation stabilizes cytoplasmic β-catenin, allowing it to translocate into the nucleus where it opens chromatin by displacing the repressor Groucho at TCF/LEF binding sites and promotes the expression of Wnt target genes (**Fig. 1A**)^24^. Genetic effects on canonical Wnt signaling are associated with the expression of complex brain traits. For example, common genetic variants associated with brain structure are enriched near Wnt pathway genes^2^, neural cells derived from individuals with neuropsychiatric disorders show alterations in expression of Wnt pathway genes^18–20^, and rare variants in Wnt pathway genes are associated with neuropsychiatric disorders, such as autism spectrum disorder (ASD)^21, 22^. Our results characterize context-specific genetic effects in hNPCs that provide novel insights into neurodevelopmental gene regulatory mechanisms underlying brain trait-associated loci.

**Figure 1:**
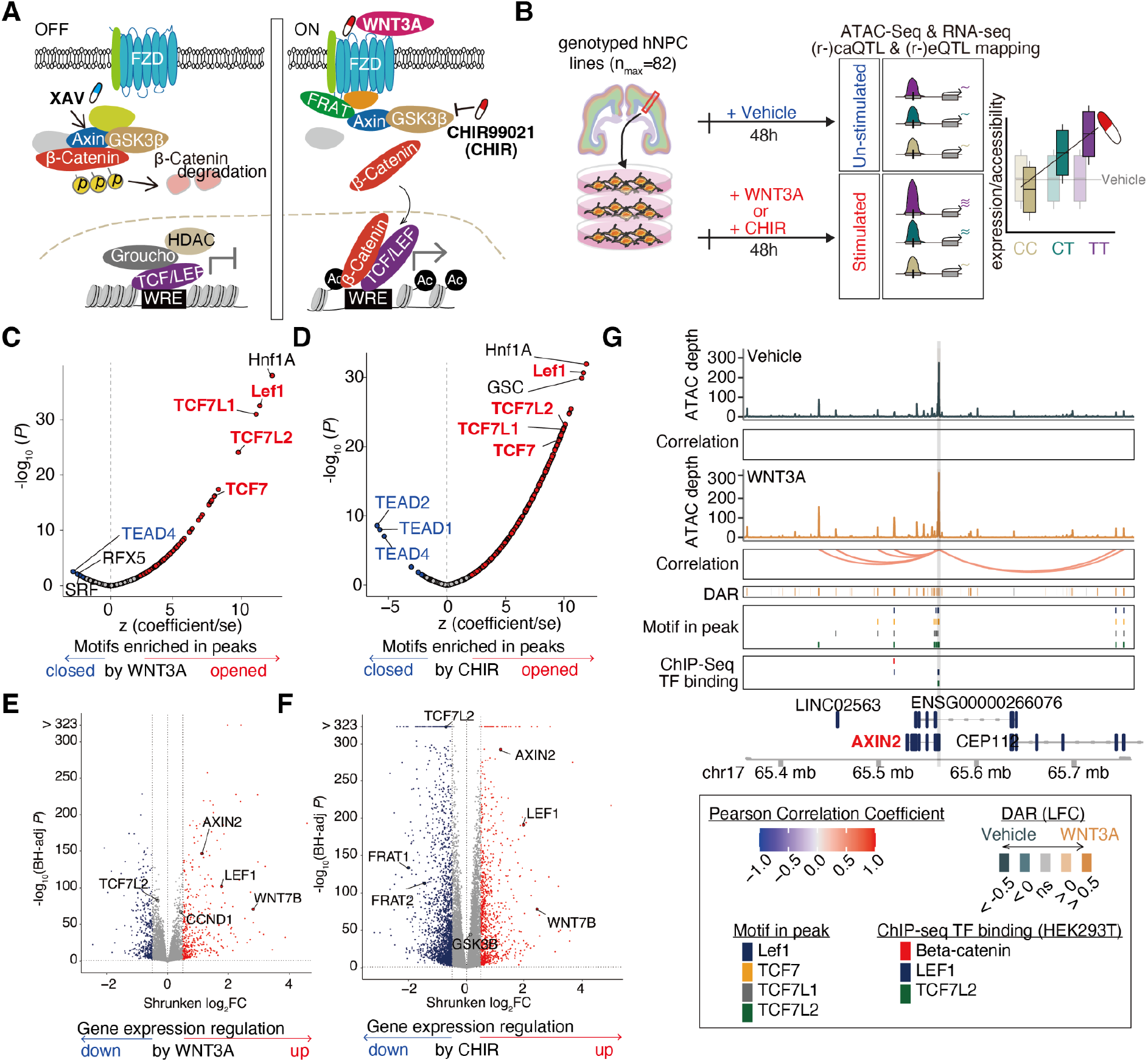
Gene regulatory changes induced by WNT stimulus. (**A**) Cartoon of the canonical WNT signaling pathway. (**B**) Schematic of study design. Enrichment of TF motifs in WNT3A-responsive (**C**) or CHIR-responsive (**D**) chromatin accessibility peaks. Z-scores reflect scaled enrichment scores (x-axis), and -log10(P-values) depict the significance of enrichment (y-axis). TFBS motifs significantly enriched in peaks opening or closing due to the stimulus are represented by red and blue points, respectively. Volcano plots show gene expression changes induced by exposure to WNT3A (**E**) or CHIR (**F**). Genes with significantly increased or decreased expression (DEGs) are represented by red and blue points, respectively. (**G**) WREs significantly correlated to AXIN2 expression during WNT3A stimulation. DAR: Differentially accessible chromatin regions measured by ATAC-seq. DARs and TSSs in the region overlap TCF/LEF motifs and Wnt-relevant TF binding from previous ChIP-seq experiments^25^.

## Results

### Wnt stimuli impact gene regulation

We interrogated gene regulation in the context of Wnt stimulation in a population of previously genotyped multi-ancestry hNPC donors that we cultured and maintained as proliferative neural progenitors^10, 26^ (**Fig. 1B, fig. S1**). To optimize activation of canonical Wnt signaling, we exposed hNPCs to various concentrations of either the WNT3A endogenous ligand, CHIR (CHIR99021, also known as CT99021), a potent GSK3β inhibitor and Wnt activator, or vehicle, for 48 hours^27^. We evaluated the effect of each stimulus on canonical Wnt-signaling using a β-catenin-responsive luciferase reporter assay^28^ and influence on hNPC proliferation using an EdU incorporation assay (**fig. S2**). Guided by the results from these assays, we selected 5nM WNT3A because it maximized both Wnt activity and proliferation. While CHIR exposure exceeding 2.5µM increased Wnt pathway activation, we selected this concentration based on its ability to maximize hNPC proliferation (**fig. S2**). Following 48h stimulation by 5nM WNT3A, 2.5µM CHIR, or vehicle, we performed ATAC-seq and RNA-seq for all samples, including 2-6 replicates for each of six randomly selected donors to evaluate technical reproducibility (**fig. S3-5**). After quality checks and selection of one technical replicate from each donor-condition pair (see Methods), we detected expression of 15,762 protein-coding genes and 7,695 lncRNAs from 242 RNA-seq samples (nvehicle=79, nWNT3A=82, nCHIR=81) and chromatin accessibility at 172,887 peaks from 222 ATAC-seq samples (nvehicle=76, nWNT3A=68, nCHIR=78).

To determine the gene regulatory impacts of Wnt stimulation on human neural progenitors, we performed differential analyses of chromatin accessibility and gene expression between WNT3A- or CHIR-stimulated conditions compared to vehicle control. Stimulation by WNT3A or CHIR revealed 21,383 unique differentially accessible chromatin regions (Wnt-responsive elements or WREs; FDR Benjamini-Hochberg-adjusted P < 0.1, |LFC| > 0.5; WNT3A vs Vehicle (62 pairs): 4,819 WREs; CHIR vs Vehicle (72 pairs): 20,179 WREs; **fig. S6A**, **table S1**). We anticipated Wnt stimulation would increase chromatin accessibility at TCF/LEF binding sites as β-Catenin displaces the chromatin condenser Groucho^24^. Consistent with these expectations, WREs opening due to Wnt stimulation were strongly enriched with TCF7, TCF7L1/2, and Lef1 motifs (**Fig 1C-D**, **table S2**). β-Catenin, Lef1, and TCF7L2 binding sites defined by ChIP-seq in HEK293T cells^25^ also overlapped WREs opened by Wnt stimulation significantly more than WREs closed by Wnt stimulation (**fig. S7**). Additional enrichment of HNF1a motifs within WREs (**Fig. 1C-D**) implies a coregulatory relationship with TCF/LEF, as has been previously described in cancer cells^29^. Interestingly, binding motifs of non-canonical Wnt signaling such as TEAD4^30^ were enriched in WREs that closed in response to Wnt stimulation (**Fig. 1C-D**), suggesting an antagonistic relationship between canonical and non-canonical WREs. These results show that Wnt stimulation in human neural progenitors modulates known downstream DNA-binding protein effectors in expected directions and defines a set of human brain-developmental WREs.

We detected a total of 3,254 unique Wnt-responsive differentially expressed genes (DEGs) across the two Wnt-stimulating conditions (DEGs, FDR-adjusted *P* < 0.1; |LFC| > 0.5; WNT3A vs Vehicle (75 pairs): 762 DEGs; CHIR vs Vehicle (74 pairs): 3,031 DEGs; **Fig. 1E-F, fig. S6B, table S3**). DEGs included known components of the Wnt pathway such as *LEF1* and *AXIN2*, confirming that Wnt stimulation leads to autoregulation of the Wnt pathway^31, 32^. Pathway enrichment analysis of DEGs showed that those upregulated in response to Wnt stimulus were over-represented in Wnt-related pathways such as “TCF dependent signaling in response to WNT” (FDR-adjusted *P* = 3.81×10^-7^ and 4.69×10^-7^, for WNT3A vs Vehicle, or CHIR vs Vehicle, respectively), as expected (**table S3-4**). The Wnt pathway is also known to increase proliferation (**fig. S2**), and indeed, upregulated DEGs were enriched in “Cell Cycle [REAC]” related genes (FDR-adjusted *P* = 7.17 x 10^-51^ and 7.97 x 10^-45^, for WNT3A vs Vehicle, or CHIR vs Vehicle, respectively)^33^. Additionally, Cyclin D1 (*CCND1*), a known target gene of WNT stimulation and a key factor regulating cell cycle progression^34^, was significantly upregulated under WNT3A stimulation (LFC = 0.44, FDR-adjusted *P* = 1.63 x 10^-67^, **Fig. 1E**).

### Wnt stimulation recruits novel regulatory elements

Activation of the Wnt-signaling pathway alters gene expression patterns that modulate NPC cellular behaviors such as proliferation and differentiation to shape brain development^35, 36^. To identify novel WREs that may impact gene expression, we estimated the correlation between chromatin accessibility and gene expression for proximal gene-peak pairs (+/-1MB from the transcription start site (TSS)) to link regulatory elements to the genes they regulate. We found that across stimulation and vehicle conditions, over 5% of peaks are significantly correlated with nearby genes and over 12% of genes are significantly correlated with nearby peaks (FDR-adjusted *P* < 0.1, **table S5-6**). On average, 83% of gene-peak pairs showed a positive correlation, supporting the idea that opening chromatin usually increases, while restricting chromatin accessibility usually decreases, the expression of target genes. Wnt stimulation revealed 12,643 enhancer-gene pairs not detected in the vehicle condition, while 6,461 were lost (**fig. S8**). For example, *AXIN2*, a gene known to be upregulated by canonical Wnt signaling across tissues^24, 37^, linked to 11 or 8 peaks each harboring TCF/LEF binding sites under WNT3A or CHIR condition, respectively, yet had no significantly correlated peak-gene links found in the vehicle condition (**Fig. 1G**). These data suggest that new regulatory elements are recruited to regulate gene expression during Wnt stimulation.

### Distinct effects of Wnt stimulation or inhibition with XAV

While both WNT3A and CHIR stimulate the Wnt pathway, they induced different effects on gene expression consistent with their distinct mechanisms of action. CHIR yielded considerably more WREs and DEGs as compared to WNT3A, suggesting that this potent small molecule inhibitor of GSK3*β* induces more gene regulatory changes as compared to the endogenous ligand at their respective concentrations. This difference possibly occurs because CHIR acts downstream of WNT3A where it may more directly affect target gene expression, though concentration differences between the two stimuli make direct comparisons difficult (**Fig. 1E-F**; **fig. S6**). The expression level of *GSK3β* is upregulated in progenitor cells stimulated with CHIR (LFC = 0.15, FDR-adjusted *P* = 4.64×10^-46^), but not WNT3A (LFC = 0, FDR-adjusted *P* = 0.97; differential impact estimated by interaction term = 9.14×10^-26^). This suggests that GSK3*β* inhibition by CHIR triggers a compensatory gene expression response that is not induced by Wnt signaling activated via the endogenous ligand (**Fig. 1A**)^38^.

To test the specificity of Wnt pathway stimulations, we compared gene expression during simultaneous activation and downstream inhibition of the pathway (WNT3A + XAV^39^) with WNT3A activation alone in 6 hNPC donor lines (**fig. S9A**). As expected, inhibition of the Wnt signaling pathway suppressed expression of genes upregulated by Wnt stimulation as compared to vehicle (r = −0.59; *P* < 1 x 10^-323^; **fig. S9B**). For example, *LEF1* expression increased in response to WNT3A stimulation, and decreased following inhibition of the WNT pathway. These opposing effects provide further support that the observed gene expression changes are caused by induction of canonical Wnt signaling.

### Wnt-responsive genes and regulatory elements contribute to inter-individual differences in brain traits

Previous studies suggest that genes related to the Wnt pathway are mutated or differentially expressed in individuals with neuropsychiatric disorders^18–22^. We sought to determine whether Wnt-responsive genes further support these disease associations by testing for enrichment of Wnt-responsive DEGs in sets of brain-related disease-associated genes using curated gene-disease information from the DisGeNET database^40^. We found enrichment of Wnt-responsive DEGs among schizophrenia and ASD risk genes, (**Fig. 2A**, **table S7**) while genes not significantly differentially expressed after Wnt stimulation did not show a detectable enrichment among brain-related disease associated genes^41^. This implies that alteration in the function of Wnt-responsive genes contributes to risk for neurodevelopmental disorders.

**Figure 2:**
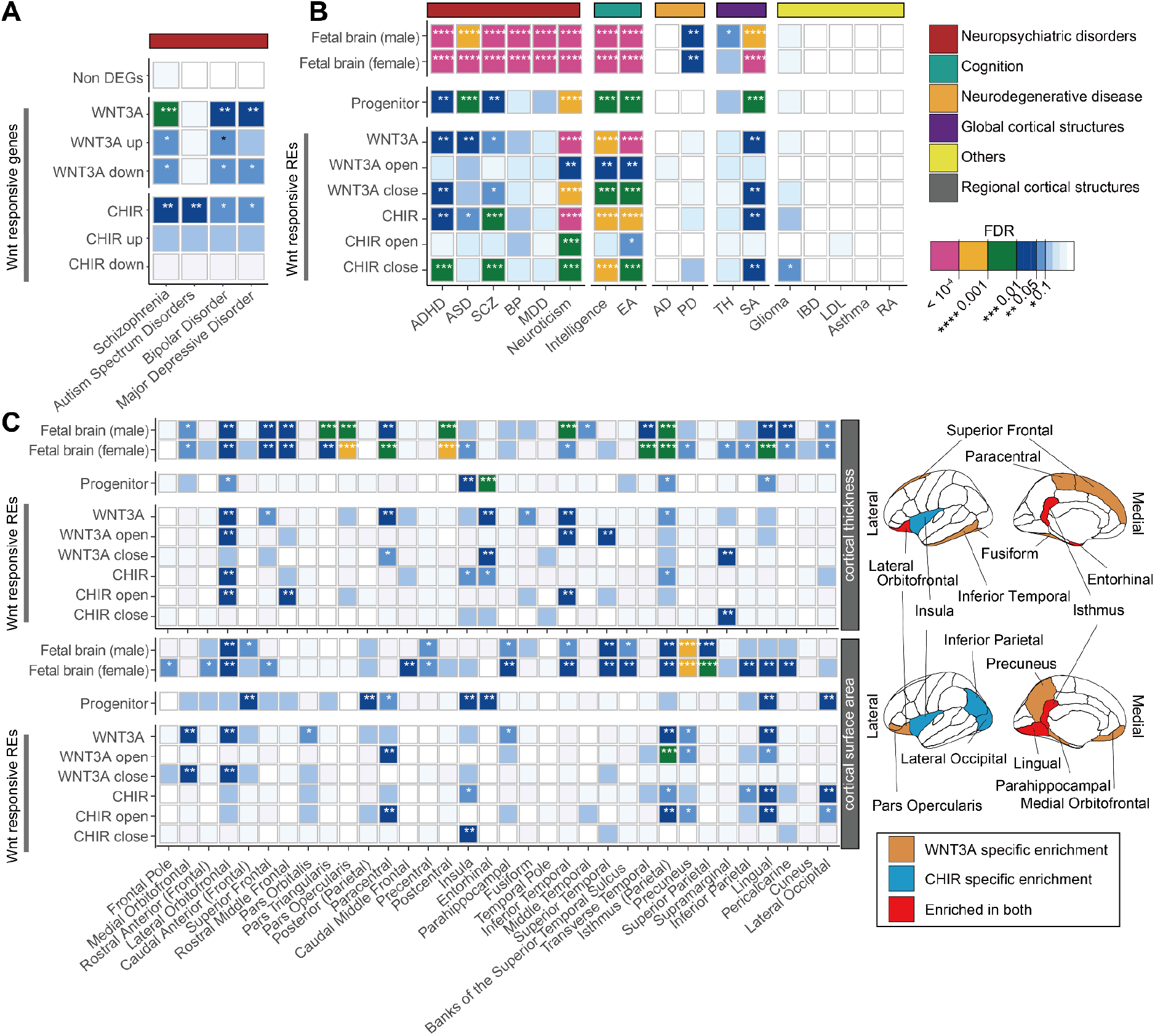
Contribution of Wnt-responsive regulatory elements to the heritability of brain traits. (**A**) Enrichment of Wnt-responsive genes within neuropsychiatric disorder risk gene sets from the DisGeNET database^40^. (**B**) Contribution of WREs to brain-related trait heritability evaluated by S-LDSC. Traits are grouped by category. (**C**) Contribution of WREs to the heritability of adult cortical thickness and cortical surface area traits across regions (left). Brain regions with significant enrichment of cortical thickness (top) or cortical surface area (bottom) traits within WREs (right). The *P* values indicated by color in (**B**-**C**) denote whether the WREs contribute significantly to SNP heritability after controlling for other annotations including elements in baseline model and/or non-WREs. * indicates enrichments with FDR < 0.1. ADHD: Attention deficit hyperactivity disorder, ASD: Autism spectrum disorder, SCZ: Schizophrenia, BP: Bipolar disorder, MDD: Major depressive disorder, EA: Educational attainment, AD: Alzheimer’s disease, PD: Parkinson’s disease, TH: Average cortical thickness, SA: Cortical surface area, IBD: Inflammatory bowel disease, LDL: low-density lipoprotein, RA: Rheumatoid arthritis (see **table S8** for references).

We next explored whether common genetic variants within WREs contribute to brain-related traits (**table S8**). By performing stratified LD score regression (S-LDSC) accounting for the baseline-model^42, 43^, we replicated previous findings that regulatory elements in fetal brain tissues or hNPCs contribute to the heritability of neuropsychiatric disorders and brain-related traits^10, 14^ (FDR-adjusted *P* < 0.1; **Fig. 2B**, **table S9**). We then applied S-LDSC to WREs and found that they contribute to the heritability of schizophrenia, ADHD, ASD, and glioma, as well as inter-individual differences in intelligence, even when controlling for the effects of non-differentially accessible peaks (**Fig. 2B**, **table S9**). We also found that common variants within WREs significantly contribute to inter-individual differences in global cortical surface area, but not cortical thickness, consistent with Wnt regulating progenitor proliferation and the predictions of the radial unit hypothesis^15, 44^. We further estimated partitioned heritability enrichment for regional cortical surface area and thickness traits. We observed regional specificity where heritability was enriched in WREs for the surface area of regions such as lingual gyrus and isthmus of the cingulate (**Fig. 2C**). Interestingly, the heritability of cortical thickness was also enriched in WREs within regions including lateral orbitofrontal and the isthmus of the cingulate. We did not detect partitioned heritability enrichment for neurodegenerative disorders (Alzheimer’s disease and Parkinson’s disease) or non-brain related traits (Irritable Bowel Disease, Low-Density Lipoprotein, Asthma, and Rheumatoid Arthritis), showing the specificity of these enrichments. In summary, common variants within Wnt-responsive genes and regulatory elements contribute to inter-individual differences in brain structure, neuropsychiatric disease risk, and cognitive ability.

### Context-specific genetic effects on chromatin accessibility and gene expression

Because Wnt-responsive gene expression and regulatory elements contribute to inter-individual differences in brain traits, we sought to identify common single nucleotide polymorphisms (SNPs) and indels affecting gene regulation during Wnt-stimulation. We mapped chromatin accessibility and expression quantitative trait loci (ca/eQTL) using stringent control for known and unknown confounding and use of a hierarchical multiple testing correction (Methods). We identified over 43,000 caQTLs (caSNP-caPeak pairs) in each condition regulating 36,423 unique caPeaks (FDR-adjusted P < 0.1; number of caQTL pairs = 43,664 Vehicle; 57,718 WNT3A; 57,581 CHIR; **Fig. 3A-B**). We also identified ∼2,000 eQTL (eSNP-eGene pairs) in each condition regulating 3,089 unique eGenes (FDR-adjusted *P* < 0.1; number of eQTL pairs = 2,025 Vehicle; 2,075 WNT3A; 1,961 CHIR) (**Fig. 3C-D**, **tables S10-11)**. The observed effect size of vehicle ca/eQTLs in this study strongly correlated with ca/eQTL effect sizes using largely overlapping hNPC samples cultured during previous studies^10, 11^, indicating our findings are highly reproducible (caQTL r = 0.93, *P* < 1 x 10^-323^; eQTL r = 0.89, *P* < 1×10^-323^; **fig. S10**). When the same SNP was identified as both a caQTL and an eQTL for a given stimulus, we observed strong positive correlation between effect sizes on chromatin accessibility and gene expression, also as found in our previous work (**fig. S11, table S17**), showing that alleles increasing chromatin accessibility generally lead to increased gene expression.

**Figure 3:**
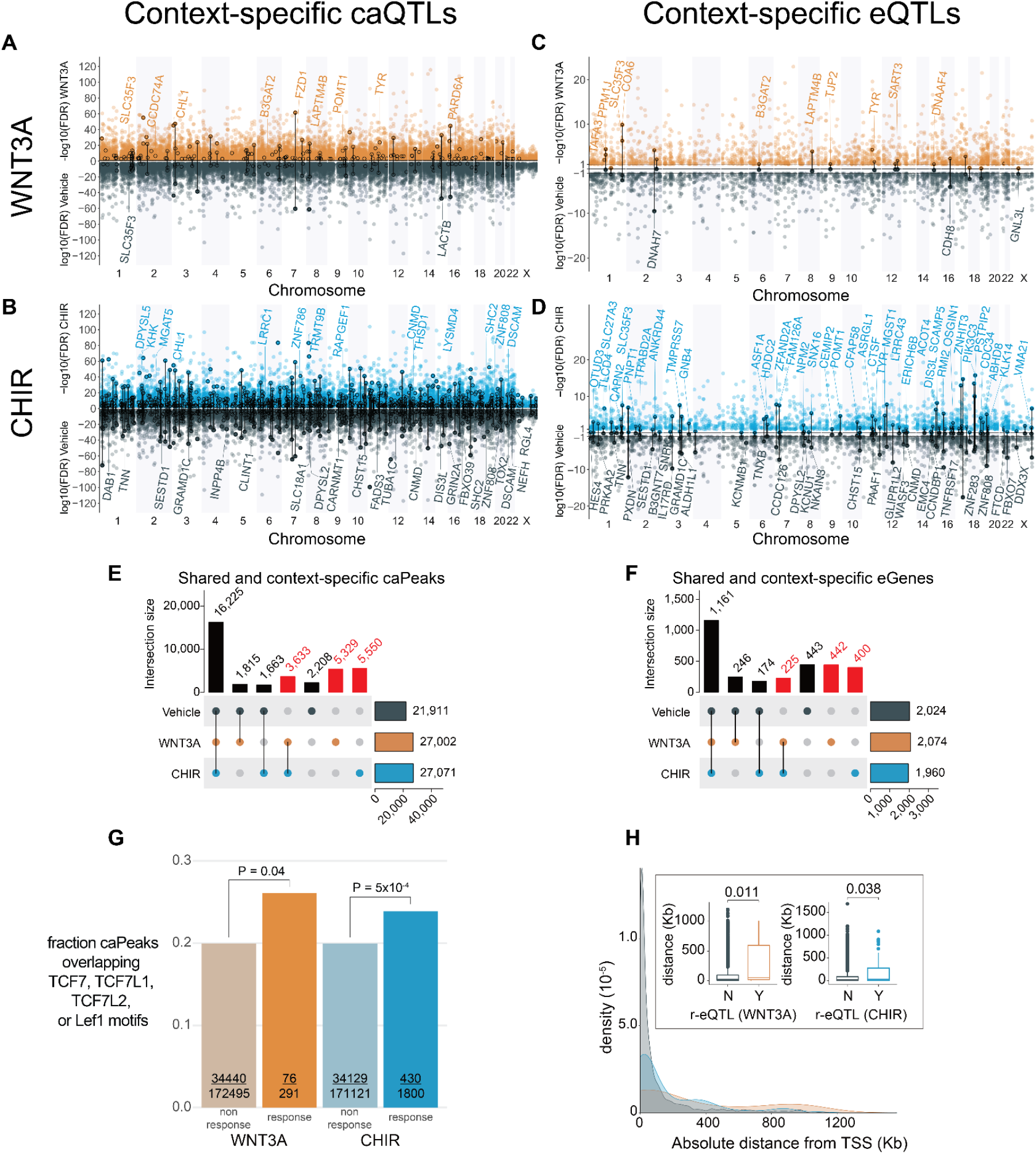
Context-specific genetic effects on chromatin accessibility and gene expression. Miami plots depict significant caQTLs (**A**, **B**) or eQTLs (**C**, **D**) detected under WNT3A (orange) (**A, C**), CHIR (blue) (**B, D**) or vehicle conditions (gray) across the genome. Circled variants denote significant genotype-by-condition interaction effects (r-QTLs). Protein-coding eGenes overlapping r-caQTLs and eGenes highlighted in subsequent figures are labeled. The number of context-specific caPeaks (**E**) or eGenes (**F**) shared across vehicle, WNT3A, and CHIR conditions. Columns labeled in red represent caPeaks or eGenes only detected in Wnt-stimulated conditions. (**G**) Fraction of response-caQTL-regulated caPeaks vs all other caPeaks containing TCF7, TCF7L1, TCF7L2, or Lef1 motifs. *P-*values denote significance of the difference in proportions of response and non-response caPeaks containing a TCF/Lef motif, evaluated by logistic regression. (**H**) Distributions of absolute genomic distances between eQTLs and their target gene’s transcriptional sites (TSS). Boxplots summarizing these distances are shown in the inset.

We observed a 66.2% increase in caPeaks and a 52.7% increase in eGenes detected in the Wnt stimulated states as compared to vehicle (**Fig. 3E-F**). The directionality and magnitude of QTL effect sizes between stimulated and unstimulated conditions were generally consistent, but nevertheless many QTLs exhibited differential effects in the stimulated condition consistent with the increase in detection of caPeaks and eGenes (**fig. S12**). These results show that Wnt stimulation reveals context-specific genetic effects on gene regulation previously undetected in unstimulated cells.

Together, context-specific caQTLs and eQTLs enable inference of enhancer priming, where a genetic variant is associated with chromatin accessibility in both unstimulated and stimulated conditions, but only leads to changes in gene expression in the stimulated condition. In this way, enhancers are primed to drive gene expression upon recruitment of additional stimulus-specific TFs. In total, we detected 397 primed regulatory elements. Primed regulatory elements were significantly enriched in a variety of genomic annotations from the fetal brain called by chromHMM, including transcription start sites, enhancers, and bivalent enhancers, but not heterochromatin, quiescent or transcribed regions (**Fig. 4A-B**). Surprisingly, we found that primed elements were not more enriched in bivalent enhancers as compared to non-primed elements. Instead, primed elements were more enriched near active transcription start sites than non-primed elements, possibly due to the greater detection of eQTLs at promoter regions. One example of a primed peak was found at a caPeak 53 kb from the TSS of CLINT1, where we detected two high LD SNPs within the peak strongly associated with chromatin accessibility in both vehicle and under CHIR stimulation, one of which disrupts the CTCF motif (**table S12**). But, this locus was only associated with gene expression under CHIR stimulation, presumably due to the recruitment of β-catenin to TCF/LEF motifs present in this peak (**Fig. 4C-E**). The CLINT1 protein interacts with clathrin to mediate endocytosis, a process important for both secretion of WNT ligands and WNT-induced accumulation of β-catenin in the nucleus^45, 46^.

**Figure 4:**
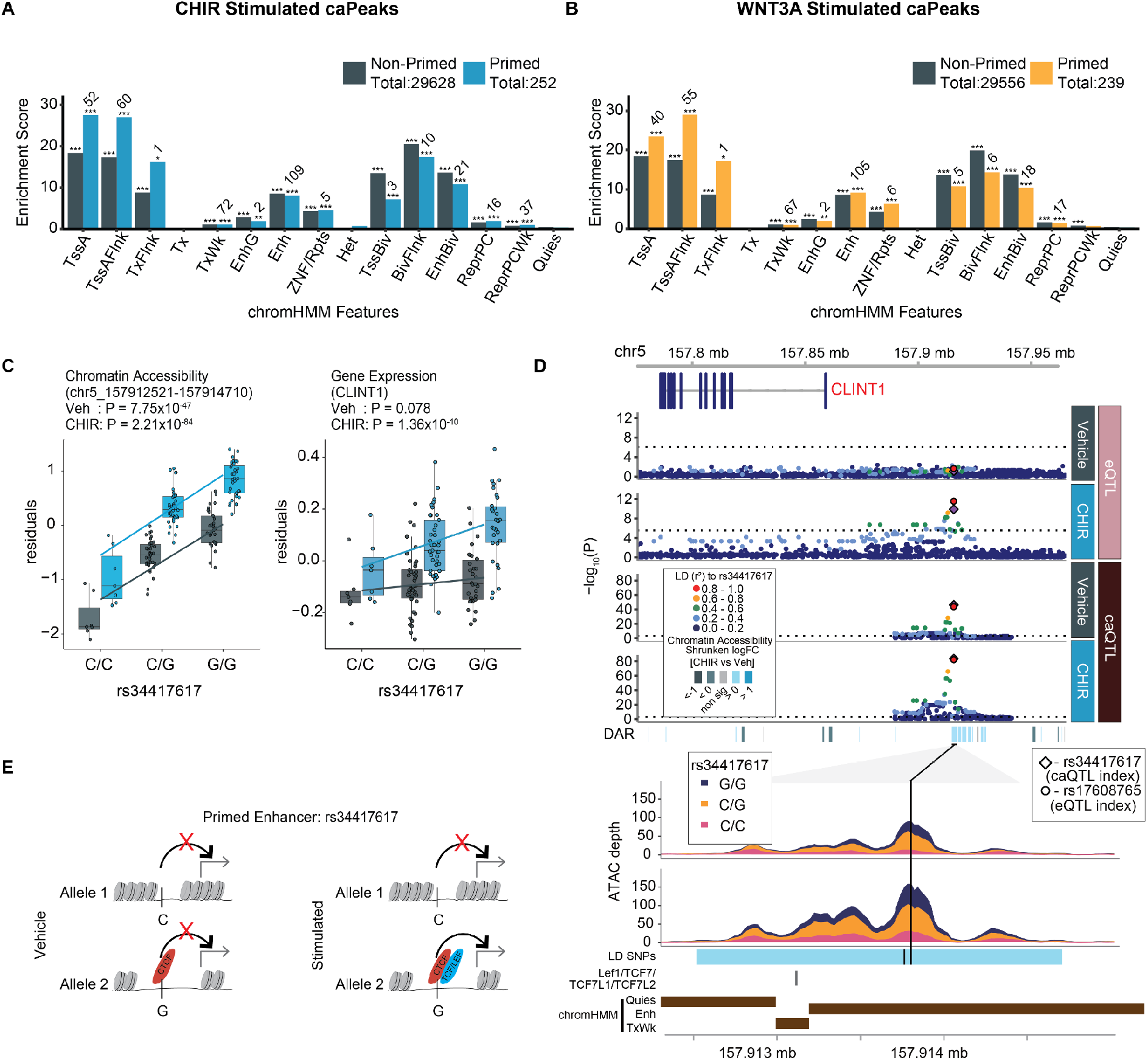
Enhancer priming identified through context-specific molecular QTLs. Enrichment of primed and non-primed CHIR (**A**) and WNT3A (**B**) caPeaks within chromHMM states defined in the fetal brain. *, **, *** indicate enrichments with *P*-values < 0.05, .01 and .001, respectively. Numeric labels indicate overlap count of a caPeak with a given annotation. (**C**) Allelic effects of rs34417617 on chromatin accessibility of WRE (chr5:157912521-157914710) (left) and *CLINT1* expression (right). (**D**) Regional association plots at the *CLINT1* locus. From top to bottom: Genomic coordinates, gene models, eQTL and caQTL *P* values for vehicle and CHIR-stimulated conditions, ATAC-seq coverage showing differential chromatin accessibility, with SNPs linked by LD (r^2^ > 0.8), TCF/LEF elements annotated, and chromHMM features. (**E**) Putative mechanism for rs34417617 regulating chromatin accessibility and gene expression in the vehicle and stimulated conditions.

Since we observed shared and distinct effects of genetic variants on gene regulation across stimulation conditions, we hypothesized two common interpretable scenarios out of many possibilities. (1) Wnt-activation increased power to detect condition-specific ca/eQTLs due to changes in accessibility or expression, or (2) Wnt activation alters the function of a genetic variant in regulating chromatin accessibility or target gene expression by revealing a differential genotypic effect as compared to vehicle. To test the latter hypothesis, we performed a genotype-by-condition interaction test for all independent QTLs, which showed the main effects on chromatin accessibility or their target eGenes, and detected 291 and 1,800 r-caQTLs, and 22 and 102 significant r-eQTLs in WNT3A, CHIR, respectively (labeled with circles in **Fig. 3A-D**, **fig S12**, **tables S13-14**). Chromatin accessibility peaks regulated by r-caQTLs were enriched in specific transcription factor binding site (TFBS) motifs as compared to all other caPeaks (**table S15**). For example, significantly more TCF7, TCF7L1/2, and Lef1 motifs were found within CHIR or WNT3A response peaks as compared to non-response peaks (**Fig. 3G**), suggesting that these Wnt-stimulation-specific TFs lead to context-specific genetic effects on chromatin accessibility. Interestingly, the transcription factor motif most significantly associated with response-caPeaks, ARID3A, is involved in neural fate specification^47, 48^, and interacts with TCF7 to co-bind regulatory elements in murine T cell progenitors^49^. Response caPeaks were enriched within several chromHMM genomic annotations including active TSS, enhancers, and bivalent enhancers. There was significantly less enrichment of response caPeaks in active TSS and enhancers as compared to non-response caPeaks, perhaps because these response caPeaks flag novel condition specific enhancers not annotated in post-mortem fetal brain tissue (**fig S13**). We also observed that r-eQTLs were more distal to the TSS of the regulated eGene as compared to non-r-eQTLs (**Fig. 3H**). This finding is consistent with the idea that context-specific regulatory elements are farther from the genes they regulate than are non-context-specific regulatory elements^50^.

### Genomic regions under strong positive selection are enriched in caPeaks

Recently, human ancestor quickly evolved regions (HAQERs) were identified describing regions of rapid sequence evolution following the divergence between humans and chimpanzees from their most recent common ancestor^51^. HAQERs are highly enriched within bivalent chromatin states in the developing human brain, sites which are thought to regulate expression of context-responsive, developmentally relevant genes^52^. We found 37 HAQERs (out of 1581 that were defined) overlapped genetically regulated caPeaks in both stimulated and unstimulated conditions, a highly significant enrichment given the size and number of these elements in the genome (**Fig 5A-B**, **table S16**). Interestingly, Wnt-stimulation led to a greater detection (60% increase) of genetic variants affecting evolutionarily relevant chromatin accessibility peaks, consistent with the idea that these highly mutated genomic elements have gained context specific function along the human lineage. One HAQER-caPeak overlap was found near PAX8 and PSD4. This peak was Wnt responsive in both WNT3A and CHIR conditions and contained a common variant that showed consistent effects on chromatin accessibility regardless of stimulation condition (**Fig 5C-D**). PAX8 expression was detected in our data across all conditions, though an eQTL was not detected for this locus to support regulation of a target gene. Another caPeak-HAQER overlap was detected at the promoter region of HAR1A/B, a region with significant sequence alterations in humans compared to other primates^53^ (**fig S14**). Only two HAQER-caPeaks also showed genetic associations with gene expression in our study: a HAQER-r-caQTL (significantly different genetic effects in vehicle and CHIR) colocalized with an eQTL for ALG1L9P in both stimulated and unstimulated conditions and a HAQER-caQTL colocalized with LINC02361 in both stimulated and unstimulated conditions. We also tested for caPeak overlap with a distinct and largely non-overlapping evolutionary annotation defined as regions highly conserved until the most recent common ancestor between humans and chimpanzees, human accelerated regions (HARs)^54^. We found 123 HARs (out of 2751 that were defined) overlapped caPeaks, yielding a highly significant enrichment both across and within stimulated and unstimulated conditions (**Fig 5B**, **table S16**). More stimulus-specific caPeaks were found overlapping HARs as compared to unstimulated caPeaks. These overlaps show that some non-coding genetic elements that underwent strong positive selection along the human lineage harbor common variation functioning during brain development with many revealed only in the context of Wnt signaling.

**Figure 5:**
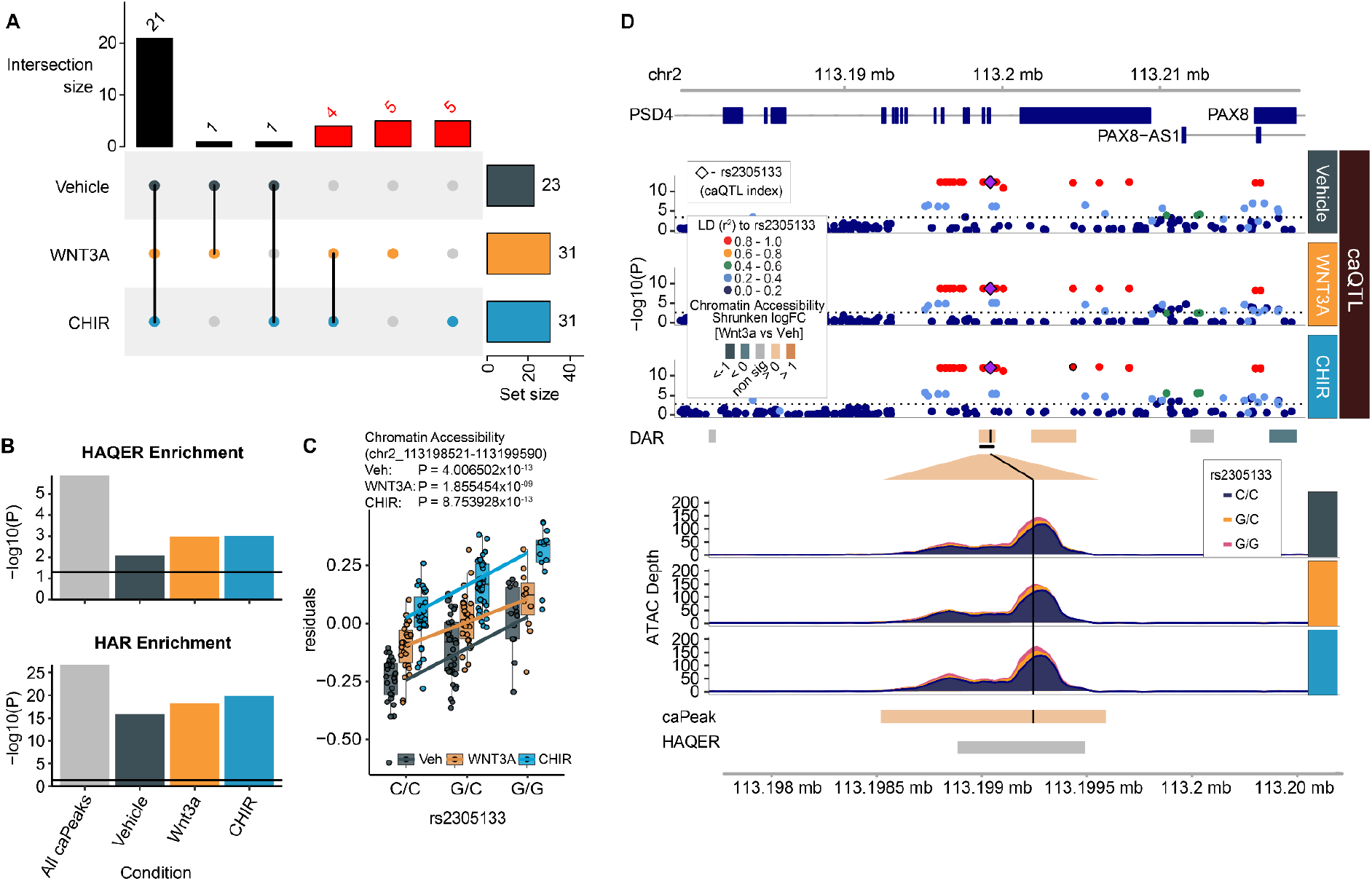
caPeaks overlap with regions under strong positive selection along the human lineage. (**A**) The number of caPeaks overlapping HAQERs across vehicle, WNT3A, and CHIR conditions. Columns labeled in red represent overlaps only detected in Wnt-stimulated conditions. (**B**) Enrichment -log10(P) values of HAQERs (top) and HARs (bottom) within unique caPeaks across all conditions and within each condition. (**C**) Allelic effects of rs2305133 on chromatin accessibility (chr2:113198521-11319959). (**D**) Regional association plots at rs2305133, the index SNP for a caPeak-HAQER overlap. From top to bottom: Genomic coordinates, gene models, caQTL P-values for vehicle, WNT3A, and CHIR-stimulated conditions, ATAC-seq coverage showing differential chromatin accessibility and HAQER location.

### Identifying context-dependent gene regulatory loci shared with brain-related GWAS traits

Partitioned heritability analysis revealed that GWAS loci associated with neuropsychiatric disorder risk and brain structures are enriched at WREs, demonstrating that these elements contribute broadly to brain phenotypes (**Fig. 2**). However, enrichments do not nominate specific genes and variants underlying these contributions. In order to identify variants and putative gene regulatory mechanisms to explain brain trait GWAS loci including brain structure, function, neuropsychiatric disorders, and cognitive ability, we examined caQTLs and eQTLs with LD-overlap to GWAS loci (**table S8**). Based on this analysis, we identified 1,684 regulatory elements and 169 genes involved in brain-traits in the vehicle condition (**Fig. 5A**). The use of stimulated conditions increased the number of brain-trait associated peaks by 72.2% and genes by 57.3% (**Fig. 5B**), demonstrating that they may explain some of the ‘missing regulation’ underlying GWAS loci. 6,965 caQTL-GWAS pairs and 1,189 eGene-GWAS pairs were unique to stimulated conditions (**Fig. 5C**, **table S18-19**). 34% of eGenes regulated by any eQTL detected in our study that overlapped GWAS loci were not previously reported as eQTLs detected in bulk postmortem human “Brain_Frontal_Cortex_BA9”^55^, representing novel overlaps specific to developing hNPCs. A subset of these eGenes (14% of all brain-trait associated eGenes detected in our study) were detected only in WNT3A- or CHIR-stimulated conditions and not in vehicle or adult bulk postmortem tissue, showing that stimulus specific eQTLs reveal novel mechanisms underlying GWAS loci undetected without stimulation (**table S19**).

We highlight two stimulus-specific colocalizations, confirmed by conditional analysis, with the r-e/ca-QTLs. First, a CHIR r-eQTL modulating expression of *ANKRD44* (rs979020-T, **Fig. 5D, fig. S15A**) colocalized with schizophrenia GWAS and the volume of the left presubiculum body hippocampal subfield^56, 57^. *ANKRD44* encodes an ankyrin repeat domain functioning as a regulatory subunit of protein phosphatase-6 (PP6), an enzyme that regulates the cell cycle and suppresses NF-kB signaling, a pathway known to engage in cross-talk with Wnt signaling^58–61^. The T allele of rs979020 is associated with increased expression of *ANKRD44*, decreased risk of schizophrenia, and reduced volume of the hippocampal presubiculum. This colocalization underscores the connection between decreased hippocampal volume and schizophrenia^62, 63^ and suggests Wnt-responsive regulation of *ANKRD44* in neural progenitors plays a role in the expression of these traits. A second example is the colocalization of a CHIR r-caQTL (rs1992311; **Fig. 5E**, **fig. S16**) with a CHIR-responsive *DPYSL5* eQTL signal and a GWAS of average thickness of the isthmus cingulate region. *DPYSL5*, also known as *CRMP5*, has shown to be a negative regulator of neural progenitor proliferation^64^. A Pou5f1::Sox2 motif is predicted to be disrupted by the caSNP in this caPeak (rs4665363-G; **fig. S16E**) which is likely modulated in the stimulation condition by TCF/LEF binding to motifs present in the same peak. This suggests that rs4665363 is a putative causal variant altering chromatin accessibility and downregulating *DPYSL5* expression, which may lead to increase of average thickness of the isthmus cingulate. We also observed stimulus-specific colocalizations supported by eCAVIAR, including *FADS3* with bipolar disorder^65^, and *ENO4* with variants associated with regional cortical surface area including insula^2^ (**fig. S18-S19**). These results highlight that fine-mapping via integrating r-QTLs and GWAS traits support putative regulatory mechanisms that impact brain-related GWAS traits.

## Discussion

In this study, we stimulated the Wnt pathway in a library of human neural progenitor cells and measured chromatin accessibility and gene expression across the genome. Wnt stimulation robustly altered chromatin accessibility and gene expression including opening of chromatin at TCF/Lef motifs, expected through β-Catenin displacing the chromatin condenser Groucho, and increased expression of known Wnt-pathway target genes including those associated with cell cycle^31^ (**Fig. 1A-F**). We defined a comprehensive set of Wnt-responsive regulatory elements present in human neural progenitor cells, and show that these elements are recruited during stimulation to regulate Wnt-responsive genes, like AXIN2 (**Fig. 1G**). WREs strongly contribute to heritability for a variety of psychiatric disorders, brain structure, and cognitive traits, implying that these brain-trait associated variants function during patterning of neural progenitors in fetal development and contribute to inter-individual differences in adult brain traits (**Fig. 2**). Inherent genetic variation in this hNPC library led to significant differences in chromatin accessibility at over 30,000 regulatory elements, and significant differences in gene expression at over 3,000 genes (**Fig. 3**). Interestingly, Wnt stimulation led to detectable impacts on regulatory elements and genes that were undetected in unstimulated states and enabled inference of gene regulatory relationships. Our results show that genetic variation has context-specific function, even within a single cell-type (**Fig. 3,4**). Some genetically influenced regulatory elements have undergone strong positive selection along the human lineage, indicating they are context-specific regulatory elements developed along the human lineage and are important for the evolution of the human brain (**Fig. 5**). In addition, genetically influenced regulatory elements and genes revealed additional mechanisms underlying GWAS signals, providing regulatory elements, genes, cell types, and cell states that impact psychiatric disorder risk and other brain-related traits (**Fig. 6**).

**Figure 6:**
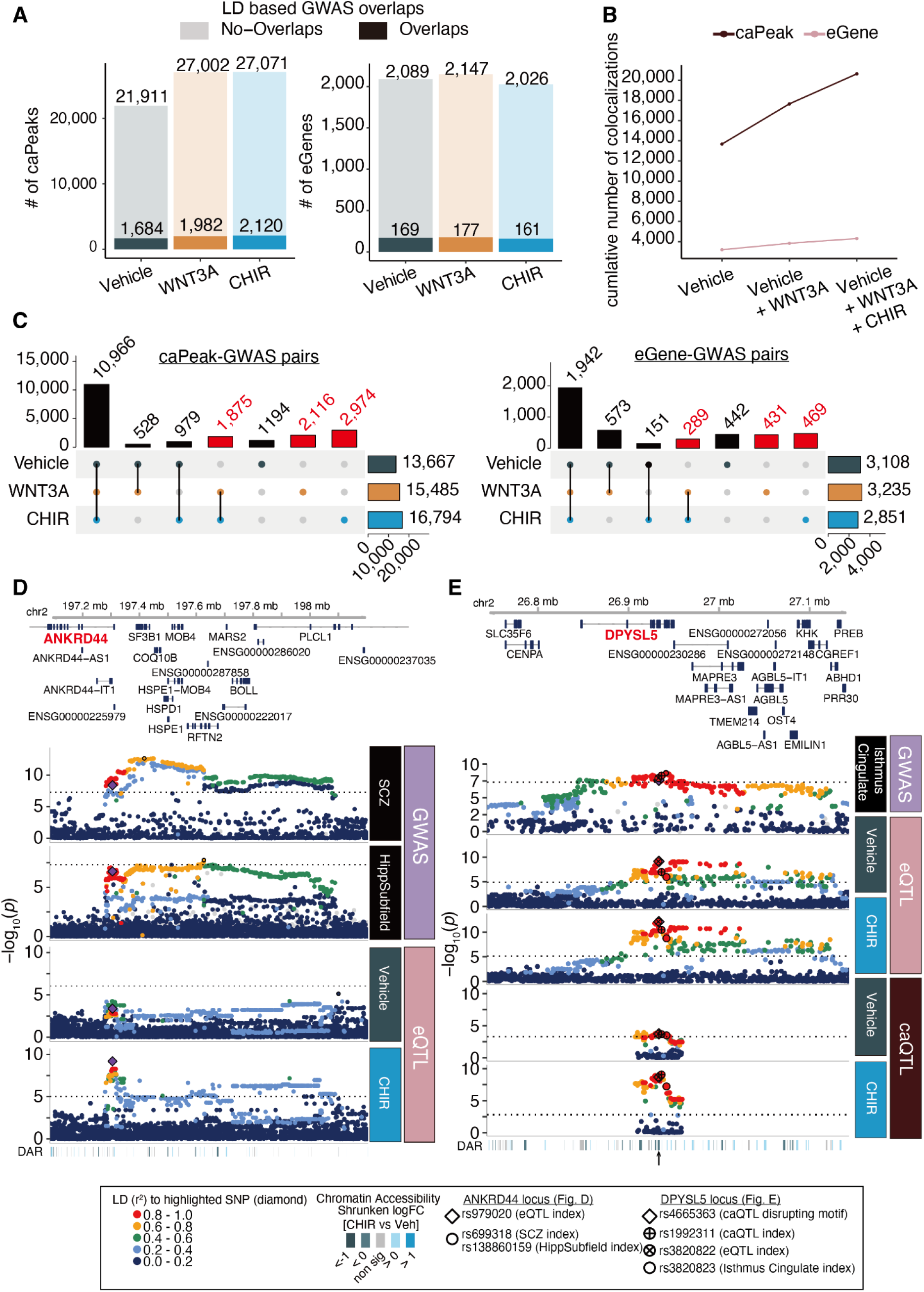
Using WNT stimulus-specific gene regulation to inform mechanisms underlying complex brain traits. (**A**) The number of caPeaks (left) or eGenes (right) overlapping brain-related GWAS loci defined by moderate LD (*r^2^* > 0.6 in either 1KG EUR population or our study) across vehicle, WNT3A and CHIR conditions. (**B**) The cumulative number of colocalized caPeaks or eGenes increases across stimulation conditions. (**C**) Shared or condition-specific caPeaks (left) or eGenes (right) colocalized with brain-related GWAS traits in each condition. Columns labeled in red indicate colocalizations only detected in Wnt-stimulated conditions. (**D**) Regional association plot depicting colocalization of schizophrenia (top panel) and the volume of a hippocampal subfield (presubiculum body, left hemisphere) GWAS with a CHIR-responsive eQTL modulating *ANKRD44* expression (rs979020-T, CHIR vs vehicle interaction FDR-adjusted *P* = 0.09). From top to bottom: Genomic coordinates and gene models, *P* values for brain-related GWAS, *P* values for condition-specific QTLs discovered in this study, and differentially accessible regions (DAR) within the locus. Differences in the patterns of association are likely due to population differences in LD between the GWAS and QTL studies. (**E**) Regional association plot depicting colocalization of average thickness of isthmus cingulate GWAS with a CHIR-responsive eQTL modulating *DPYSL5* expression and a CHIR-responsive caQTL (rs1992311, interaction FDR-adjusted *P* = 0.041; chr2:26932281-26934470, in an intron of *DPYSL5*), arranged as in (**D**).

Our results show that the function of some genetic variants are dependent on environmental stimuli. New genetic variant function can be revealed through stimulation, where no detectable effect on gene regulation is observed in unstimulated states (**Fig. 3, 6D**). Conversely, genetic variant function can be hidden during stimulation but only revealed in unstimulated states (**Fig. 3**). Another possibility is that genetic variants can have an effect in both unstimulated and stimulated states, but a stronger effect in one condition showing that the stimulus modulates a variant’s effect (**Fig. 3,6E**). All scenarios indicate that stimulation alters the function of genetic variation and are detectable through interaction analyses, but future development of statistical tools that can separate these possibilities will aid biological interpretations.

Most research focusing on understanding the regulatory function of non-coding genetic variation relies on bulk adult post-mortem tissue which cannot respond to external stimuli and lacks cell type specificity. Our study addresses this limitation and reveals novel mechanisms underlying GWAS traits where eGenes regulated by eQTLs overlapping GWAS loci were undetected by bulk adult post-mortem eQTLs^55^14% of eGene-GWAS overlaps only found following Wnt stimulation were novel compared to frontal cortex GTEx eQTLs. These observations underscore that consideration of stimulus, cell-type, and developmental time illuminates some of the ‘missing regulation’ linking genetic variants to the expression of these traits. Because our study design employed prenatally-derived primary human neural progenitor cells stimulated using a potent developmental signaling pathway involved in cell proliferation and brain patterning, our results imply that genetic variants exert context-dependent effects during early neurodevelopment that can lead to differences in adult brain and behavioral traits.

Paired chromatin accessibility and gene expression QTLs allowed us to detect 397 gene regulatory relationships indicative of enhancer priming, where genetic effects on chromatin accessibility are present in both unstimulated and stimulated conditions, but genetic effects on gene expression require a stimulus^9^ (**Fig. 4**). Bivalent elements are defined as those with both active and repressive chromatin marks. These elements have been posited to poise regulatory elements for quick activation of target genes, though little support for this function has been found^52, 66^. We found that while primed enhancers were enriched in regions with bivalent chromatin marks defined in the developing human brain, they were less enriched than non-primed elements. While this could be due to differences in eQTL detection power, another possibility is that priming and poising are not necessarily overlapping mechanisms. We hypothesize that bivalent enhancers tend to regulate gene expression that establishes commitment to cell fate decisions and operate on a relatively slower time-scale that requires editing of chromatin modifications^52^. In contrast, primed “enhancers” may regulate genes in a relatively faster and more plastic manner dependent on binding of stimulus specific transcription factors and do not require both active and repressive chromatin marks.

We noted significant overlap between our caPeaks and regions of rapid evolution in human ancestors defined by cross-species sequence comparisons (HAQERs/HARs)^51, 54^(**Fig. 5A,B**). This overlap provides further evidence that at least some HAQERs/HARs are indeed functional stimulus-specific regulatory elements active during brain development. HAQERs are highly mutable and enriched for common genetic variants associated with a variety of neuropsychiatric disorders, further supporting that genetic variation within evolutionarily relevant hot-spots influences the expression of complex brain traits^67^. Here we show that some of the common genetic variants within both HAQERs and HARs can influence their inferred activity, and in turn may influence traits related to brain development. We speculate that HAQERs and HARs represent functional regulatory elements that affect hNPC fate decisions potentially driving human-specific cortical expansion.

caQTL and eQTL mapping revealed novel effects of common genetic variants undetected in unstimulated states. While sample size limits QTL discovery, and especially response-QTL power, we discovered thousands of Wnt stimulus-specific caPeaks and eGenes. Genetic effects detected under unstimulated conditions replicated those found in previous studies of unstimulated hNPC QTLs, underscoring the robustness of in vitro QTL study design. We focused on studying genetic variation during stimulation of the well-studied and developmentally important Wnt signaling pathway, because alterations in this pathway have been associated with risk for neuropsychiatric disorders, all brains are exposed to this stimulus during development, activators and inhibitors are known, and the downstream effectors of the signaling pathway on gene expression are known^2, 15–23^. Expansion of this approach in large populations of primary human neural progenitor cells or induced pluripotent stem cells can investigate other gene-by-environment interactions in a dish. Specifically, similar study designs evaluating stimulation for different durations, stimulation of additional signaling pathways relevant to brain development^68^, exposures to environmental insults^69^, responses to clinically useful drugs^70^ or modulation of neuronal activity^71^, may reveal additional genetic effects that are masked in QTL studies conducted in bulk post-mortem tissue.

## Methods

### hNPC culture

#### Ethics statement for human tissue-derived cell-lines

This study followed IRB regulations to derive human NPC cell-lines from prenatal tissue collected at the UCLA Gene and Cell Therapy facility following voluntary termination of pregnancy.

#### Generation of hNPC lines

Fetal brain tissue visually consistent with dorsal telencephalon morphology (flat and sheet-like) was collected at 14-21 gestation weeks from presumed neurotypical donors to derive primary human NPCs as previously described^10, 11, 26^. To summarize, tissue was dissociated into single cells which were cultured as neurospheres before transfer to fibronectin (Sigma F1141) and Poly-L-Ornithine (Sigma P3655) coated plates. Neuronal differentiation was suppressed to maintain NPCs in a proliferative state by following previously established culture methods^10^. After 2-3 passages, NPC lines were cryopreserved and transferred to UNC Chapel Hill. NPC media: Neurobasal A (Life Technologies 10888-022) supplemented with 100 μg ml^−1^ primocin (Invivogen ant-pm-2), 10% BIT 9500 (Stemcell Technologies 09500), 1% glutamax (100x; Life Technologies 35050061), 1 μg ml^−1^ heparin (Sigma-Aldrich H3393-10KU), 20 ng ml^−1^ EGF/FGF (Life Technologies PHG0313/PHG0023), 2 ng ml^−1^ LIF (Life Technologies PHC9481) and 10 ng ml^−1^ PDGF (Life Technologies PHG1034).

#### hNPC cell culture and WNT-stimulation

We cultured cryopreserved hNPCs in batches of 8 cell lines per week, pseudorandomizing each experimental group for biological variables (sex and donor gestation week), and technical variables (passage number). Following a two-week expansion period, we plated 400k NPCs per well of a 6-well plate for each cell-line. Next, we exposed each well to either vehicle (Neurobasal A media supplemented with PBS+0.1%BSA and DMSO), 5nM Wnt3a in PBS+0.1% BSA, or 2.5uM CHIR in DMSO for 48h. All exposures were prepared such that equal volumes were applied for each stimulated condition, including a “balancer” solution composed of culture media, PBS+0.1% BSA, DMSO, and water in order to standardize diluents across all exposures. The well-position of each exposure was rotated every week of the experiment to minimize the potential effects of plate position. After 48h exposure to stimuli or vehicle, cells were lifted with Accutase (Thermo Fisher Scientific A1110501) for preparation of ATAC-seq and RNA-seq libraries. We generated 2-6 separately cultured replicates for each of 6 randomly selected donors to measure technical variation across the experiment. We utilized cell culture media, growth factors, additives, and WNT-stimulating exposures from the same manufacturer lots across the entire experiment, whenever possible, to mitigate potential batch effects. When it was not possible to use the same lot of reagent, this information was recorded. One individual (JMV) performed all hNPC cultures and exposures to stimuli to minimize variance in handling effects. The investigators were not explicitly blinded to the donor during cell culture or library preparation. However, the investigators did not have knowledge of donor genotype when performing cell culture or library preparation. Regular pre-assay screening detected no mycoplasma contamination (ATCC 30-1012K).

### hNPC Wnt activity luciferase assays

Canonical Wnt pathway signaling activity was measured via a luciferase reporter assay (**fig. S2A-C**). After two weeks of hNPC culture expansion as described above, we lifted cells with 0.05% trypsin (Gibco 25300062) and plated 10k cells counted by hemocytometer per well of 96-well plates (Corning 07-200-91). Cells were transduced with 10uL lentivirus carrying BAR:Luciferase and Tk:Renilla constructs 24 hours after plating. Plasmids were generous gifts from the lab of Ben Major^72^ and lentivirus was generated as previously described^70^. After a two-day incubation, cells were exposed with either vehicle, CHIR, or WNT3A as described above. 48 hours later, cell lysates were collected and luminescence measured using the Dual-Glo luciferase system (Promega E2920) on the GloMax Discover plate reader (Promega). β-Catenin induced BAR:Luciferase signal was normalized by Renilla luminescence from the constitutively active Tk promoter.

### hNPC proliferation assays

After two weeks of hNPC culture expansion as described above, we lifted cells with 0.05% trypsin (Gibco 25300062), and plated 12.5k cells counted by hemocytometer per well of 96-well plates (Corning 3610). We generated 4 technical replicate wells for each concentration of Wnt-activating stimulus or vehicle, and 8 technical replicate wells for vehicle exposures per hNPC donor cell-line, distributing replicates across each culture plate to mitigate potential artifacts introduced by plate position. One day later, vehicle or Wnt activating stimuli were applied, as described above. 46 hours later, we exposed cells with 10 μM EdU with a 10% media addition for 2 hours. hNPCs were collected and fixed via a 10-minute incubation in 4% paraformaldehyde in phosphate-buffered saline. On these cells, we labeled total DNA content with FxCycle Far Red dye (Thermo Fisher Scientific F10347) and performed Click-iT EdU-incorporation assays (Thermo Fisher Scientific C10337) according to the manufacturer’s protocols. We quantified EdU and FxCycle signals using the Attune NxT 96-well Flow Cytometer (Thermo Fisher Scientific) (**fig. S2D-F**), or via high content imaging on the Nikon Eclipse Ti2 microscope followed by manual quantification with Fiji (**fig. S2G-H**). Flow cytometry data was processed with FlowJo v10.7.1 and cell-cycle populations defined by automated gating with FlowDensity software^73^. Detailed methods for analysis of flow cytometry can be found in previous work^70^.

### ATAC-seq Library preparation

Cells were lifted and counted to isolate 50k cells per sample to be used as input for the Omni-ATAC protocol for library preparation^74^. To summarize, 50k nuclei from fresh NPCs were counted via hemocytometer and tagmented using TDE1 enzyme (Illumina 20034198) and tagmentation buffer composed of 20mM Tris-HCl pH 7.6, 10mM MgCl_2_, and 20% Dimethyl Formamide. All libraries were PCR amplified (NEB M0544S) for 5 cycles, and then each library was further amplified for a sample-specific additional number of cycles (average 2 additional cycles across all samples) determined by a qPCR side reaction to avoid overamplification. Nextera-compatible unique dual-indexing primers barcoded each sample during amplification (Illumina 20027213). We purified libraries to remove primer dimers and eliminate fragments over 1000bp (Roche 07983298001). We validated ATAC-seq library quality before sequencing for a subset of samples by observing appropriate nucleosomal banding patterns via capillary gel electrophoresis on an Agilent 4150 TapeStation system (Agilent 5067-5584). We then pooled ∼70 ATAC-seq libraries with unique barcodes in each of four pools, and sent pooled libraries for multiplexed sequencing. One individual (BDL) generated all ATAC-seq library preparations in order to minimize batch effects introduced by handling variance.

### RNA-seq Library Preparation

Total RNA was isolated from each hNPC line and condition using all cells remaining (average 1.1M cells per sample) after removing 50k cells for ATAC-seq library preparation as described above. For each experimental week, hNPCs were lifted with Accutase (Thermo Fisher Scientific A1110501), lysed, and stored in TriZol (Thermo 15-596-026) at −80C for subsequent purification of total RNA using RNeasy Mini Kits and on-column DNase digestion (Qiagen 74106, 79256). All RNA extractions were performed by the same individual (JTM) to minimize potential batch effects introduced by handling. We quantified RNA isolates using fluorescence-based Quant-iT RNA assay kits (Thermo Q33140). Mean RNA integrity number (RIN) across all samples was 9.86 (SD = 0.39). Library preparation was performed using a stranded total RNA kit (KAPA KR0934) after samples were depleted of ribosomal RNA (KAPA KR1151).

### Sequencing of ATAC-seq and RNA-seq libraries

ATAC-seq libraries were sequenced at the NYGC to an average read depth of 49M (SD = 13.6M) paired-end reads (2×100bp) per sample on the Illumina Novaseq platform. Total RNA-seq libraries were sequenced at the NYGC to an average sequencing depth of 55M (SD = 13M) paired-end reads (2×100bp) per sample on the Illumina Novaseq platform.

### Genotyping and imputation

hNPC genotypes were obtained from genomic DNA isolated with DNeasy Blood and Tissue Kit (QIAGEN 69504) using Illumina’s HumanOmni2.5 platform. Additional variants were imputed using the TOPMed freeze 5 reference panel^75^ with minimac4 software^76^ on the University of Michigan Imputation server. We performed quality control, pre-processing, and filtering of SNPs with PLINK v1.9^77^ as previously described^10^ based on Hardy-Weinberg equilibrium, minor allele frequency, individual missing genotype rates, and variant missing genotype rate (plink --hwe 1e-6 --maf 0.01 --mind 0.1 --geno 0.05).

### ATAC-seq data preprocessing

FastQC (v0.11.9)^78^ and MultiQC (v1.7)^79^ software performed quality control for all ATAC-seq libraries before and after trimming sequencing adapters with BBMap (v38.98)^80^. We then mapped ATAC-seq reads to the hg38 reference genome with the Burrows-Wheeler Alignment tool (BWA-MEM v0.7.17)^81^. We then refined alignments to minimize mapping bias at sites where ATAC-seq reads overlapping bi-allelic SNPs using imputed genotypes by re-mapping and removing duplicate reads with WASP (v0.3.4)^82^. Samtools (v1.16)^83^ then removed unmapped or mitochondrial reads and bedtools (v2.3)^84^ removed reads mapped to genomic blacklist regions as defined by ENCODE^85^. ATAC-seq read metrics during and following preprocessing were calculated using picard (v2.21.7)^86^ and ataqv software (v1.0.0)^87^ (**fig. S4**). We evaluated cross-sample contamination using verifyBAMID^88^, and omitted samples with FREEMIX or CHIPMIX scores greater than 0.02. We also omitted samples with low transcriptional start site enrichment (TSSe < 5) and aberrant short read/mononucleosomal read ratios (S/M <1 or S/M > 7). In total, 51 samples were omitted by these criteria. Principal component analysis (PCA) on ATAC-seq samples separated samples by stimulus condition, and not by sex or other biological or technical variables in the first two principal components (PCs) of variance (**fig. S5**). Sex and Donor ID, which captures cell-line intrinsic effects for each hNPC showed some correlation with other PCs, leading us to include these variables in QTL models described below and use residualized data following regression of PCs 1-10.

### Chromatin Accessibility Peak Calling

After selecting a single ATAC-seq replicate for each donor-condition pair based first on whether stimulated conditions and vehicle were cultured in the same plate, and subsequently on on QC metrics, we called chromatin accessibility peaks from ATAC-seq reads using all samples excluding technical replicates with CSAW’s (v1.28)^89^ *windowCounts* function with the options (ext = *mean length of fragments*, filter = *5 * sample number* spacing = *10*, param = pe.param) where pe.param were *max.frag = 1500, pe=’both’, minq=20*. Windows within 100bp were merged with the *mergeWindows* function. Read counts were normalized accounting for GC-content via conditional quantile normalization with cqn software^90^.

### Differential chromatin accessibility analysis

We ran DEseq2 for the read counts described above to identify peaks with differential accessibility due to stimulus condition, called WREs. The comparisons were performed as paired tests within donor, hence, the model was set to *Peak Accessibility* ∼ *Donor ID* + *stimulus condition*, where Donor ID and stimulus condition were factor variables. We repeated this analysis for each WNT stimulus condition vs vehicle pairs (e.g. WNT3A vs Vehicle and CHIR vs Vehicle) or WNT3A+XAV vs WNT3A. FDR-adjusted *P* < 0.1 was used as the significance threshold.

### RNA-seq data pre-processing and analysis

Similarly to the preprocessing of ATAC-seq data described above, we initially screened RNA-sequencing libraries using FastQC and MultiQC. We then trimmed sequencing adapters and mapped reads to genes as annotated by Ensembl v104 for the hg38 reference genome using the STAR aligner (v2.7.7a)^91^. We filtered samples with (1) low RIN score (<7), (2) low unique mapped rate (< 80%), (3) high mismatched rate (>50%), (4) high multi-mapping rates (>8%), (5) high duplicated read rates (> 30%), or (6) FREEMIX or CHIPMIX scores from verifyBAMID greater than 0.02. QC filtering by these criteria removed 27 samples, and retained a total of 242 samples. Transcripts were collapsed using collapse_annotation.py (https://github.com/broadinstitute/gtex-pipeline/tree/master/gene_model) into a single gene model and gene-level reads were summarized for each sample using featureCounts from Rsubread (v2.8.2)^92^. As with the ATAC-seq samples, we performed PCA and found that the RNA-seq samples clustered according to stimulus condition and not sex, RIN, or other variables in PC1 vs PC2 plots (**fig S5**). Again, some PCs correlated with hNPC donor sex and RIN, indicating that we should include these variables in the QTL models described below for data corrected for PCs 1-10. Significantly higher pair-wise correlations within RNA-seq sample technical replicates generated for the same hNPC donor and condition (“within donor”) compared to correlation across distinct donors (Evaluated by the Welch two sample t-test) supports the technical reproducibility of the RNA-seq dataset (**fig. S3**).

After selecting a single RNA-seq replicate for each donor-condition pair based first on whether stimulated conditions and vehicle were cultured in the same plate, and subsequently on QC metrics, DESeq2^93^ was used to assess differential gene expression from RNA-seq reads mapped to protein-coding genes or lncRNA where at least 1% of samples in either condition show at least 10 normalized counts. We evaluated the model *RNAseq Count ∼ RIN + Donor ID + stimulus condition* to identify DEGs. In this model, we represented RIN as a numeric variable and Donor ID and stimulus condition as factor variables. Shrunken log2 fold change (LFC)^94^ was used for estimating dispersions and FDR-adjusted *P-*value < 0.1 was used as significance threshold.

### Pathway enrichment analysis on differential expressed genes

To determine pathways enriched in DEGs, we first obtained up-/ down-regulated genes in WNT stimulus condition as compared to vehicle (shrunken LFC > 0, < 0, respectively; and FDR-adjusted *P* < 0.1). All genes included in DEG analysis were used as the background. For both DEG and background genes, we restricted the analysis to protein-coding genes located outside the MHC region (chr6:28,510,120-33,480,577). Analysis was run by g:COSt from g:profiler^95^ for KEGG and REACTOME pathways^96, 97^. FDR-adjusted *P* < 0.1 was used as the statistical significance threshold.

### Motif Enrichment Analysis

We tested differential transcription factor (TF) motif enrichment within WREs, primed-caPeaks, or response-caPeaks for 841 predicted human TFs in JASPAR 2022^98^ core database (taxonomic group = vertebrates), using data downloaded from http://expdata.cmmt.ubc.ca/JASPAR/downloads/UCSC_tracks/2022/hg38/ (data access date: May 4th 2022). We restricted analysis to TFBS motifs within conserved regions, defined by 100-way phastCons scores > 0.4 downloaded from UCSC Genome Browser in regions ≥ 20bp^99^. For WREs, estimated TFBS motif enrichment within the top 2,000 chromatin accessibility peaks representing chromatin regions gained or lost in each stimulus condition (|LogFC| at FDR-adj *P* < 0.1 as compared to vehicle) to avoid bias introduced by the different number WREs across conditions. We ran a logistic regression to identify TF motifs observed more often in opened vs. closed peaks or response vs. non-response peaks while accounting for differences in peak width and percentage of peaks in conserved regions using the following model: *glm(TFBS ∼ peaktype + peadkwidth + conservedbppercent, family =’binomial’),* where *TFBS* is a binary outcome that indicates whether each differentially accessible peak overlaps with TF binding sites, *peaktype* indicates open/closed WREs, or response/non-response caPeaks for each stimulus condition, *peadkwidth* indicates window size of peak and *conservedbppercent* indicates percentage of conserved regions (*conserved bp* / *peak width*). We considered estimated effects of *peaktype* on TFBS motif presence with FDR-adjusted *P* < 0.1 as statistically significant.

### Correlation between chromatin accessibility and gene expression

We estimated Pearson’s correlation between chromatin accessibility and gene expression using variance stabilizing transformation (VST) normalized ATAC-seq and RNA-seq read counts residualized by 10 global chromatin peak or gene expression PCs, respectively. For each condition, we tested all peaks located within 1MB from the transcription starting site (TSS) of each gene. FDR-adjusted *P* < 0.1 was used for the statistical significance threshold.

### Disease enrichment analysis using differentially expressed genes

We performed disease enrichment analysis for each DEG category (upregulated/downregulated/all-regulated/non-DEGs) using disease_enrichment() function implemented in disgenet2r^40^. Disease enrichments referenced “CURATED” genes from 4,254 diseases from the DisGeNET database^40^, and we classified results with FDR-adjusted *P* < 0.1 generated by the disease_enrichment() function as significant. Due to limited space, we only presented diseases if disease_semantic_type is either “Mental”, “ Behavioral Dysfunction”, “Individual Behavior” or “Mental Process” in **Fig. 2**. Full lists are provided in **table S8**.

### Partitioning GWAS heritability by WREs

We first assessed the contribution of regulatory elements including tissue-type-specific or cell-type-specific regulatory annotations to the overall heritability of brain-related and other traits (**table S8**). Analyses were performed using Stratified LD Score regression (S-LDSC)^43, 100^ (v1.0.0) including the baseline-LD model (v1.2). Annotations included (1) Active enhancer or promoter states in fetal brains (female/male). Active enhancer and promoter regions were defined based on chromatin states predicted by chromHMM^101^ and included: ‘active transcription start site’, ‘flanking active TSS’, ’genic enhancers’, and ’enhancers’ in the core 15-state model (https://egg2.wustl.edu/roadmap/web_portal/chr_state_learning.html). (2) Progenitor-specific chromatin accessible regions were defined as logFC > 0 and FDR-adjusted *P* < 0.1 as a comparison to differentiated neuronal cells. ATAC-seq data were obtained from our previous work^10^. (3) Open or closed chromatin accessible regions by stimulus condition compared to vehicle conditions were defined by logFC > 0 or logFC < 0, FDR-adjusted *P* < 0.1 and identified in the current study as WREs. For (3), we included non-WREs (FDR-adjusted P > 0.1) in the model to test the specificity of WREs. We obtained publicly available GWAS summary statistics for testing partitioned heritability. The details of GWAS are provided in **table S8**. For each GWAS summary statistics, SNPs were filtered to those found in HapMap3^102^ using munge_sumstats.py provided by LDSC. The pre-computed LD scores for the European population from the 1,000 Genome Project Phase 3 (1KG)^103^ were downloaded from https://data.broadinstitute.org/alkesgroup/LDSCORE/eur_w_ld_chr.tar.bz2. Enrichment was assessed using *P*-values calculated by the annotation’s standardized effect size (tau) that was conditioned on other annotations included in the model. FDR-adjusted *P* < 0.1 was used for statistical significance threshold.

### caQTL and response-caQTL mapping

caQTL analyses within each condition (Vehicle, WNT3A, CHIR) were performed using the joint model implemented in RASQUAL^104^ which integrates both between-individual signals and allele-specific signals within individuals. Previously described cqn normalization factors were used as sample specific offsets. Covariates were included consisting of 10 count-based principal components and 10 multidimensional scaling (MDS) genotype components to reduce the confounding effects of technical factors and population structure, respectively. Variants were tested according to the following criteria (1) in a given accessible region or within +/- 25kb (2) minor allele frequency (MAF) ≥ 1% (3) Hardy-Weinberg equilibrium < 0.000001 (4) imputation quality ≥ 0.3 (5) at least two individuals have minor homozygous allele or heterozygous allele. Multiple testing correction followed a hierarchical correction procedure^11, 105^ consisting of (1) adjusting nominal *P*-values of all SNPs for each peak separately using the eigenMT method^106^ (2) BH procedure was then applied to these locally adjusted *P*-values to determine globally adjusted *P*-values lower than 0.1 (3) To determine other independent SNPs for each peak, the maximum nominal *P*-value from step 1 corresponding to a globally adjusted *P*-value of 0.1 was used as a significance threshold. Due to limitations of RASQUAL, conditional analysis, where the top caSNP was controlled for in the association model to find additional independent signals, was not possible. To determine signals most likely to be distinct from the lead for each peak a LD threshold of < 0.2 was used iteratively until no additional significant caQTL remained. Response caQTL interaction analyses between vehicle and each WNT-stimulus condition (WNT3a or CHIR) used the R package lme4 to evaluate the following linear mixed model for all index caQTL with main effects (n_caSNP_ = 136,163 (WNT3A_Vehicle pairs), 136,641 (CHIR_Vehicle pairs)):

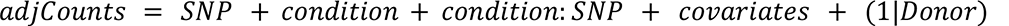

where condition = 0 or 1 (Veh or WNT activators, respectively). The first 10 count-based PCs were used as covariates. We compared the fit of the model above with and without the condition:SNP interaction term to assess significance. BH was then used to adjust for multiple testing (FDR-adjusted *P* < 0.1).

### cis-eQTL and response-eQTL mapping

Cis-eQTL analyses within each condition (Vehicle, WNT3A, CHIR) were performed using a linear mixed model implemented in limix_qtl^107^ (https://github.com/single-cell-genetics/limix_qtl), an optimized version of limix^108^. The following genes and SNPs were tested: *Gene*: (1) at least 1% of samples in the condition have at least ≥ 10 normalized counts; (2) protein-coding gene or lncRNA; *SNP*: (1) located in gene body or within +/- 1MB from gene body. (2) minor allele frequency (MAF) ≥ 1% (3) Hardy-Weinberg equilibrium < 0.000001, (4) imputation quality ≥ 0.3, (5) at least two individuals have minor homozygous allele or heterozygous allele).

We used the following model.

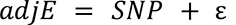

where *adj*E is VST-normalized gene read count and further corrected with 10 principal components across the gene expression matrix, that were chosen to maximize the number of eQTLs (**fig. S20**), SNP is a genotype (0/1/2), *ɛ* is an error term with cov(*ɛ*) = (σ^2^ _u_*u_k_* + σ^2^_e_*I*) where *u_K_* is the kinship matrix, σ^2^_u_ Is the variance attributable to genetic relatedness, and σ^2^_e_ is the variance attributable to random noise. The kinship matrix was generated by pcrelate() function in GENESIS (v2.14.1)^109^ with 10 PCs to control ancestry population structure.

Gene-level multiple corrections were done for each gene by permutation test implemented in limix_qtl, then applied BH-FDR for global correction. eSNP-eGene pairs with FDR-adjusted *P* < 0.1 were considered as statistically significant eQTLs. Those eGenes were further tested for independent signals through conditional testing, whereby the index eSNP is included as a covariate in the association model. For visualization purposes, we set the significance line threshold for loci with no significant eQTL in the condition as follows. First, we obtained the maximum permutation test *P*-value satisfying FDR-adjusted *P* < 0.1 (max_permP). Then, we estimated the median of the maximum raw *P*-value less than max_permP across eGene as the significant line threshold.

Response eQTL interaction analyses between vehicle and each WNT-stimulus condition (WNT3A or CHIR), were also performed by limix_qtl for all index eQTLs with main effects (n _eSNP_ = 2,966 (WNT3A_Vehicle pairs), 2,906 (CHIR_Vehicle pairs)). The interaction term for the model

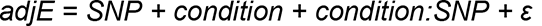

was tested to identify interaction effects between SNP and condition where condition = 0 or 1 (Vehicle or WNT activators, respectively), and *ɛ* is defined as above. We used BH to adjust multiple testing (FDR-adjusted P < 0.1).

### Effects of ca/eQTLs on transcription factor binding sites

We used motifBreakR (v.2.6.1)^110^ to assess the impact of genetic variants within peaks on TF binding motifs surrounding significant caSNP-caPeak associations (parameter setting, threshold of 1×10^-4^)^111^ (**table S12**). Annotated motifs (839 total TF motifs) from JASPAR2022 vertebrate were used in MotifDb (v1.37.1)^112^. Relative entropy (parameter setting method, ‘ic’) for both reference and alternative allele was calculated and only TFBSs strongly affected by the SNPs (parameter setting effect, ‘strong’) were retained.

### Determining Primed Enhancer Candidates

caPeaks sites were assessed for patterns indicative of priming or stimulus-specific signals within each Vehicle-WNT stimulus pair. caPeak and eGene overlaps were determined where a caPeak was within 1Mb of an eGene and at least 1 significant SNP in both datasets were in LD r^2^ ≥ 0.8 (**table S17**). Priming signals were called based on (1) significant caPeaks common to both Vehicle and WNT stimulus condition (2) only those which overlapped an eGene signal (3) significant eGene specific to WNT stimulus. Stimulus-specific signals were assessed following the same criteria but filtering to significant caPeaks specific to the WNT stimulus.

### Genomic Feature Enrichment Analysis

Primed and non-primed caPeaks were evaluated for overlaps within each Vehicle-WNT stimulus pair for overlaps with the 15 core chromHMM states defined in fetal brain male (E081)^113^. All unique caPeaks across both WNT stimulated conditions were subset into response and non-response sets and overlapped with the same chromHMM states. HAQERs (1581 total) and HARs (2751 total after liftover from hg17 to hg38) were also assessed for overlaps with genetically regulated caPeaks within each condition^51, 54^. Significance of enrichment of overlaps with previously annotated genomic features within each of these datasets were then assessed using a binomial test^114^. Neighborhood enrichment scores were calculated using a previously defined method^113^. The differences in enrichment between two annotations was calculated with Fisher’s exact test.

### Shared eSNPs/caSNPs and GWAS SNPs

We assessed colocalization between causal eca/e-QTL variants and GWAS signals in brain-related traits (**tables S18 and S19**) through a three-step process. First, we extracted GWAS SNPs (*P* < 5 x 10^-8^) that are located within 1MB from ca/eQTLs. Second, we selected GWAS-QTL pairs for which index SNPs are in LD (*r^2^* > 0.6) in either population (1KG EUR or samples used in this study). To consider the possibility of undefined secondary GWAS signals, we also selected GWAS-QTL pairs for which at least one non-index but genome-wide (GW) significant (*P* < 5 x 10^-8^) GWAS SNP was in LD with an eQTL index SNP. We tested two colocalization approaches and reported if either approach suggested colocalization. As the first approach, we estimated residual GWAS association statistics after conditioning ca/eQTL index SNPs. We performed approximate conditional analysis using default GCTA settings on variants within 1Mb of those variants^115, 116^. We used a subset of 40,000 European-ancestry UKBB participants for the LD reference panel (https://www.ukbiobank.ac.uk/). We excluded variants from GCTA output if the frequency of effect allele differed by >0.2 between UKBB and GWAS summary results, and masked approximate conditional results if variants exceeded a collinearity threshold of 0.9 with the ca/eQTL index SNP. The second approach was to use eCAVIAR^117^. We calculated SNP-level colocalization posterior probability (CLPP) for each SNP in a locus that passed the previous two steps. CLPP was estimated by eCAVIAR^117^ for SNP at P < 0.05 in both GWAS and QTL. We considered any locus with CLPP > 1% and *r^2^* between causal SNP and at least one of GW SNPs in the region > 0.8 as evidence of colocalization in this approach. False positives for QTL condition-specificity could occur due to insufficient power limited by sample size. Thus, we repeated eCAVIAR to salvage potentially shared eGene/caPeak-GWAS pairs that failed to meet FDR-adjusted significance thresholds (P < 0.1), but passed the raw significance threshold (P < 10^-6^) for QTL mapping under a particular condition, and showed shared caPeak/eGene-GWAS pairs under a separate condition.

## **Acknowledgements**

This study was supported by grants from the National Institute of Mental Health (NIMH) (R01MH118349, R01MH120125, R01MH121433). JMV and BDL were supported in part by NIH T32 training grants (T32GM135123 and T32GM067553, respectively). We gratefully acknowledge technical support from UNC research Core facilities which are supported by University Cancer Research Fund Comprehensive Cancer Center Core Support grant (P30-CA016086). The UNC Flow Cytometry Core Facility is supported in part by North Carolina Biotech Center Institutional Support Grant 2017-IDG-1025 and NIH 1UM2AI30836-01. The UNC High Throughput Sequencing Facility receives support from the UNC Center for Mental Health and Susceptibility grant (P30-ES010126). We thank Adriana Beltran and Sarahi Gabriela Molina at the UNC Stem Cell Core Facility for access to instruments. We thank Mauro Calabrese for generous access to the TapeStation instrument for sequencing library QC.

## Author contributions

BDL, JMV, and JLS designed the study. JMV maintained hNPC cultures and applied stimuli. BDL and JMW performed cellular assays. BDL generated ATAC-seq libraries. JTM isolated RNA samples. BDL and JMV processed and prepared ATAC-seq and RNA-seq data. MIL advised on statistical methods. NM, BDL, JMV and MLB performed QC and analyses on sequencing data. KAB ran conditional analyses for GWAS colocalization. DL and NA provided previous caQTLs and eQTLs, respectively, for comparison with results reported here. MJZ and KLM contributed comments and discussion to guide study design and analyses. NM, BDL, JMV, and JLS prepared the initial draft of the manuscript. All authors read and approved the final manuscript.

## Data and materials availability

All code used in analyses and data including full summary statistics except that which is provided in supplementary tables will be available upon publication at https://bitbucket.org/steinlabunc/wnt-rqtls/. We posted the current manuscript on bioRxiv (https://doi.org/10.1101/2023.02.07.527357).

## Supplementary Materials

Supplemental Figs. S1 to S20 Tables S1 to S19

