## Supplemental File for "Wnt activity reveals context-specific genetic effects on gene regulation in neural progenitors"

### Supplementary Information

**Supplementary Figures** **3**
- Supplementary Figure 1| Multidimensional scaling analysis of genotype data 3
- Supplementary Figure 2| Effects of CHIR and WNT3A on proliferation and Wnt activity 4
- Supplementary Figure 3| Technical reproducibility of RNA-seq and ATAC-seq 6
- Supplementary Figure 4| ATAC-seq Quality Control 7
- Supplementary Figure 5| Global gene expression and chromatin accessibility patterns 8
- Supplementary Figure 6| Number of differentially accessible regions and differentially expressed genes 9
- Supplementary Figure 7| Wnt-pathway related transcription factor binding sites are enriched in upregulated WREs 10
- Supplementary Figure 8| Stimulus-specific regulatory element-gene correlation 11
- Supplementary Figure 9| Opposing gene regulation by WNT pathway inhibition 12
- Supplementary Figure 10| ca/eQTL effect sizes compared to previous data 13
- Supplementary Figure 11| eQTL-caQTL overlaps 14
- Supplementary Figure 12| Effect size differences across conditions 15
- Supplementary Figure 13| Response and Non−Response caPeaks enrichment within chromHMM Annotations 16
- Supplementary Figure 14| Context-specific caPeak overlaps HAQER near HAR1A/B 17
- Supplementary Figure 15| Colocalization of ANKRD44 r-eQTL, schizophrenia, and the volume of a hippocampal subfield (presubiculum body) 18
- Supplementary Figure 16| Colocalization of an r-caQTL in DPYSL5 region and average thickness of the isthmus Cingulate 19
- Supplementary Figure 17| Shared and context-specific QTL GWAS overlaps confirmed by eCAVIAR 20
- Supplementary Figure 18| Stimulus-specific GWAS colocalization of FADS3 and Bipolar disorder 21
- Supplementary Figure 19| Stimulus-specific ENO4 eQTL colocalizing with regional cortical surface area GWAS 22
- Supplementary Figure 20| Optimizing control for known and unknown technical confounding in eQTL 23

**Supplementary Tables** **24**
- Supplementary Table 1| Differentially accessible peaks 24
- Supplementary Table 2| TF motif enrichment analysis 24
- Supplementary Table 3| Differentially expressed genes 24
- Supplementary Table 4| Pathway enrichment analysis 25
- Supplementary Table 5| Summary of peak-gene correlation 26
- Supplementary Table 6| Peak-gene pairs 26
- Supplementary Table 7| Disease enrichment analysis results(DEG) 26
- Supplementary Table 8| Summary of GWASs used in this study 27
- Supplementary Table 9| Partitioned heritability analysis results 27
- Supplementary Table 10| List of caQTL - caPeak 28
- Supplementary Table 11| List of eQTL - eGene 29
- Supplementary Table 12| caSNP TF motif disruption (in peak) 30

Supplementary Table 13| List of response-caQTLs 30
Supplementary Table 14| List of response-eQTLs 31
Supplementary Table 15| TFBS motif enrichment in response-caPeaks 32
Supplementary Table 16| caQTL Regional Overlaps 32
Supplementary Table 17| caQTL-eQTL colocalizations 33
Supplementary Table 18| GWAS colocalization results (caQTL) 33
Supplementary Table 19| GWAS colocalization results (eQTL) 34
**Supplemental Information References 35**

### Supplementary Figures

#### Supplementary Figure 1| Multidimensional scaling analysis of genotype data

Multi-dimensional scaling (MDS) plot shows the first two components of genetic similarity for all HapMap populations and donors in this study, allowing inference of genetic ancestry. Multiple ancestries of donors are present in this population.

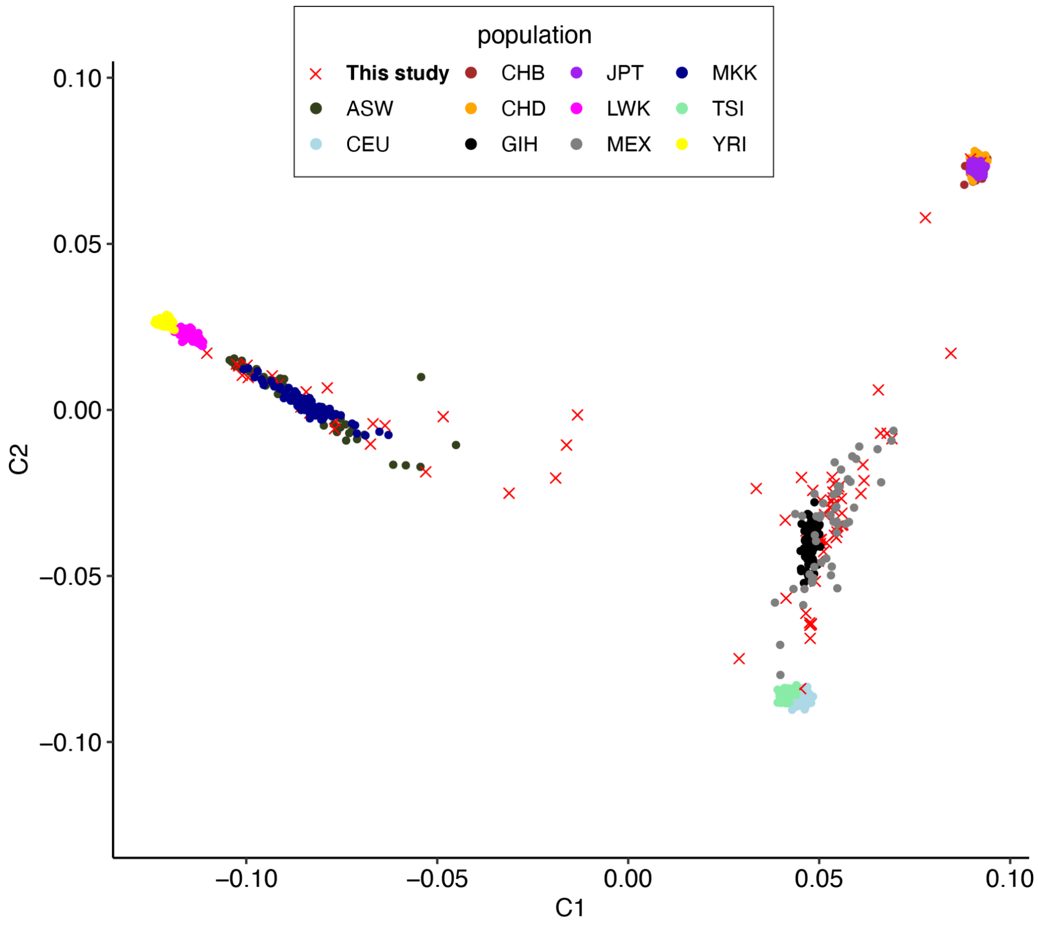

### Supplementary Figure 2| Effects of CHIR and WNT3A on proliferation and Wnt activity

Cellular assays show that WNT3A and CHIR stimulation increase canonical Wnt pathway signaling and hNPC proliferation. Diagram of luciferase reporter assay measuring β-catenin mediated Wnt pathway activation by CHIR or WNT3A (**A**). Effects of 48h CHIR (**B**) or WNT3A (**C**) exposure on Wnt pathway activation, reported as the log of luciferase luminescence (activated by Wnt stimulation) normalized by renilla luminescence (from a constitutively active reporter transgene). Representative flow cytometry scatter plots from proliferation assays depict newly synthesized DNA from a 2h hr EdU pulse vs total DNA content following 48h vehicle (left) or 2.5uM CHIR (right) exposure (**D**). Percentage of cells in S-phase (%EdU+) exposed to vehicle or increasing doses of CHIR for 48h as measured by flow cytometry in a subset of cell-lines (**E**), and in all cell-lines used for ATAC-seq and RNA-seq in this study (**F**). Representative immunocytochemistry images (100x magnification) from proliferation assay of hNPCs following 48h vehicle (left) and 5nM WNT3A (right) exposure (**G**). Green (GFP) labels cells in S-phase during the EdU pulse, and blue (DAPI) stains all nuclei. Percentage of cells in S-phase (%EdU+) exposed to vehicle or increasing doses of WNT3A for 48h as measured by flow cytometry in a subset of cell-lines (**H**). CHIR and WNT3A concentrations (boxed) that maximize Wnt activation and proliferative responses were used in this study for ca/eQTL mapping.

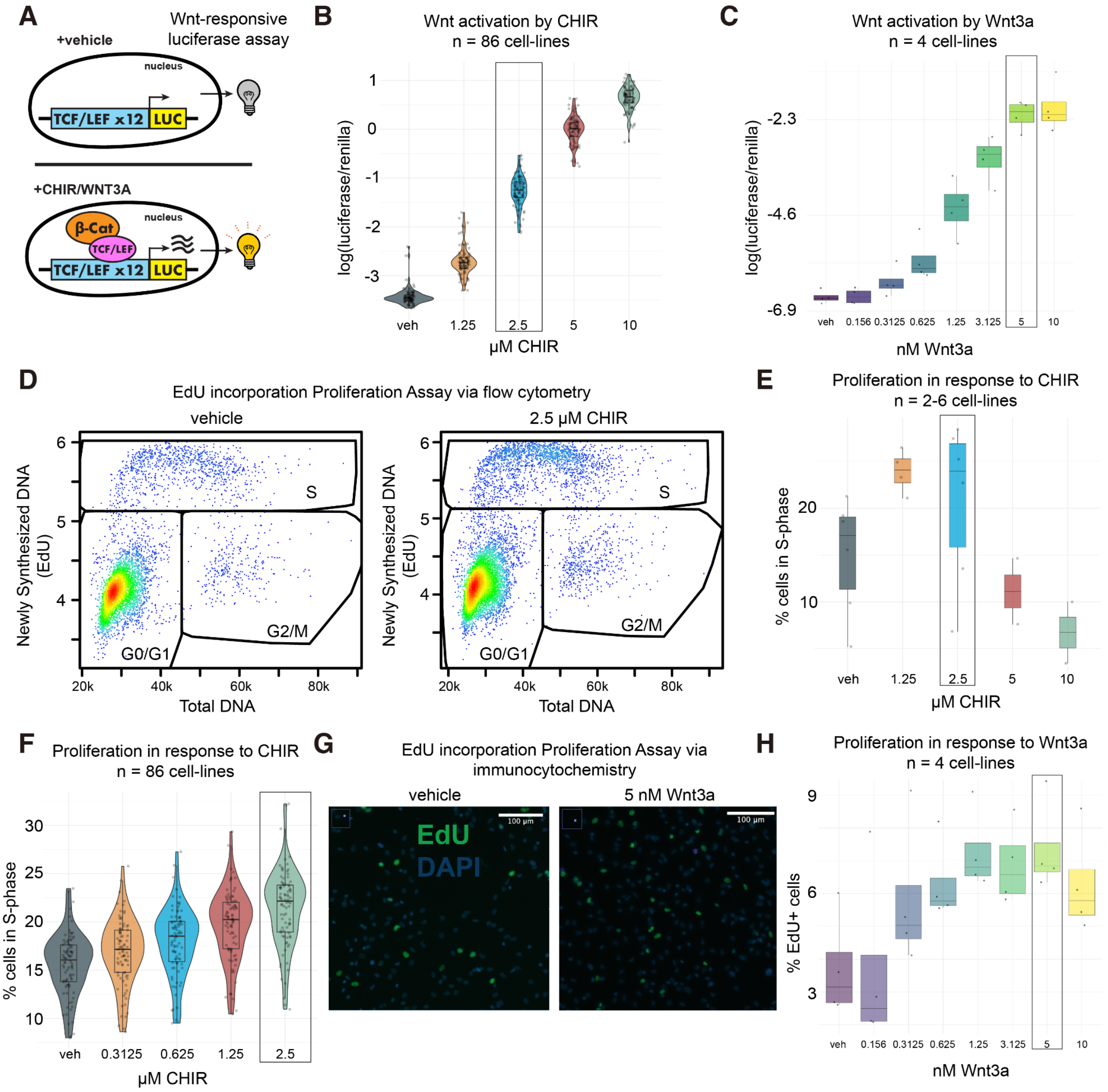

A
Wnt-responsive luciferase assay
+vehicle
nucleus
TCF/LEF x12
LUC
+CHIR/WNT3A
β-Cat
TCF/LEF
B
Wnt activation by CHIR
n = 86 cell-lines
log(luciferase/renilla)
µM CHIR
veh
1.25
2.5
5
10
C
Wnt activation by Wnt3a
n = 4 cell-lines
log(luciferase/renilla)
-2.3
-4.6
-6.9
nM Wnt3a
veh
0.156
0.3125
0.625
1.25
3.125
5
10
D
EdU incorporation Proliferation Assay via flow cytometry
vehicle
2.5 µM CHIR
Newly Synthesized DNA (EdU)
Total DNA
S
G2/M
G0/G1
20k
40k
60k
80k
E
Proliferation in response to CHIR
n = 2-6 cell-lines
% cells in S-phase
µM CHIR
F
Proliferation in response to CHIR
n = 86 cell-lines
% cells in S-phase
µM CHIR
veh
0.3125
0.625
1.25
2.5
G
EdU incorporation Proliferation Assay via immunocytochemistry
vehicle
5 nM Wnt3a
EdU
DAPI
100 µm
H
Proliferation in response to Wnt3a
n = 4 cell-lines
% EdU+ cells
nM Wnt3a

### Supplementary Figure 3| Technical reproducibility of RNA-seq and ATAC-seq

Pairwise correlations normalized sequencing counts from technical replicates across open chromatin peaks measured by ATAC-seq (**A**) or genes measured by RNA-seq (**B**). Higher correlations for within donor vs across donor pairs indicates robust reproducibility of measurements for a given donor and condition. Violin plots represent the distribution of Pearson correlation coefficients calculated between a pair of genotypically distinct donors (“across donor”) or between two technical replicates of the same donor cultured at different times (“within donor”). Top and bottom horizontal lines within violin plots represent the interquartile range of the data, and the middle bar represents the median. P-values report significant differences between the across vs. within donor correlations following Fisher’s Z transformation and evaluated with the Welch two sample t-test. The total number of cell-lines and pairwise correlations (n) for each experimental condition is reported along the x-axis.

A

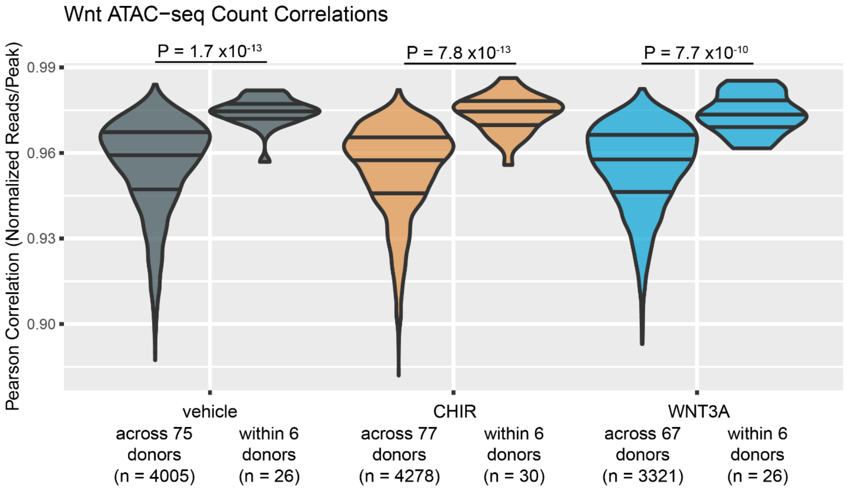


B

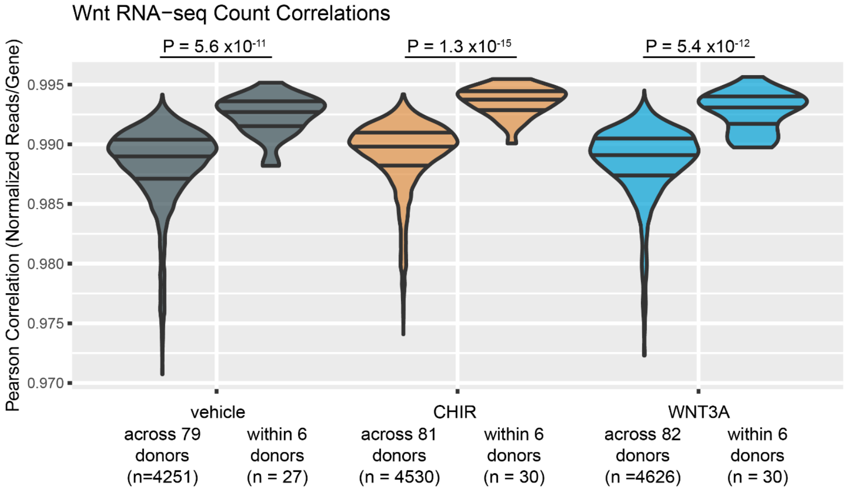

### Supplementary Figure 4| ATAC-seq Quality Control

Total ATAC-seq reads (**A**), unique non mitochondrial reads (**B**), transcription start site (TSS) enrichment (**C**), total read duplication rate (**D**), mitochondrial read duplication rate (**E**), and fraction of reads within chromatin accessibility peaks (FRiP) (**F**) across stimulus conditions. Mean insert size distributions in base pairs across all samples colored by stimulus condition (**G**) exhibit a nucleosomal phasing pattern; right: insert size distribution plotted on log- scale. Mean TSS enrichments (> 7) and FRiP scores (>0.3) exceed the "ideal" metrics for ATAC-seq libraries defined by the ENCODE project[1]. Together, these measurements validate the quality of ATAC-seq samples used in this study.

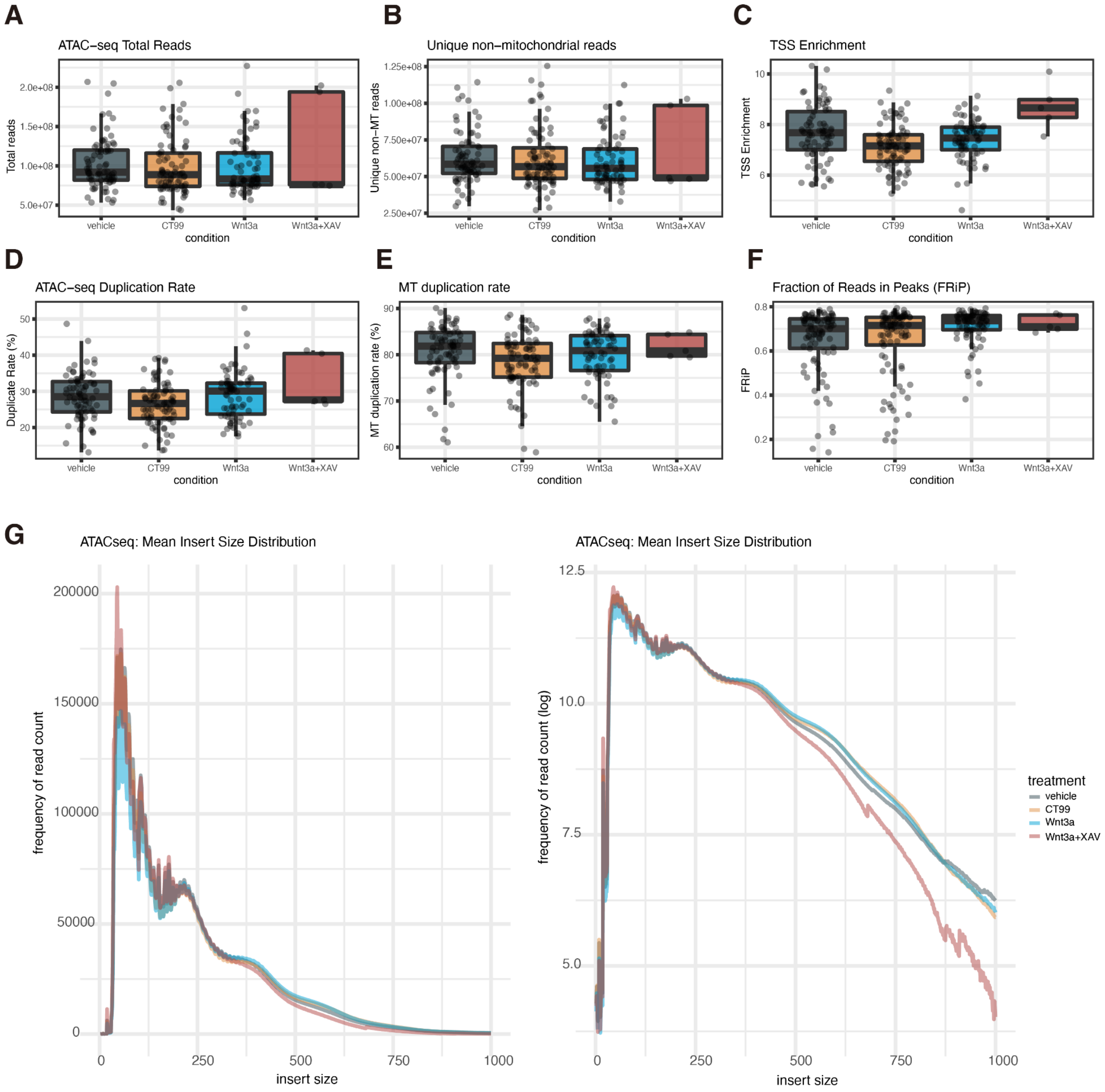

### Supplementary Figure 5| Global gene expression and chromatin accessibility patterns

Principal component analysis (PCA) of ATAC-seq (**A, C**) and RNA-seq (**B, D**) data after batch correction for technical variables included in differential accessibility or expression models, respectively. Samples are labeled by condition (**A**, **B**), or sex (**C**, **D**). Variance in global gene expression and chromatin accessibility profiles across the first two PCs is driven by stimulation condition, but not by sex. Correlation matrix of ATAC-seq (**E**) or RNA-seq (**F**) principal components 1-10 with technical (RIN) and biological variables (donor and sex). We performed linear regression to remove effects of PCs 1-10 and the residualized sequencing count data was used as QTL model input to account for measured and unmeasured confounding.

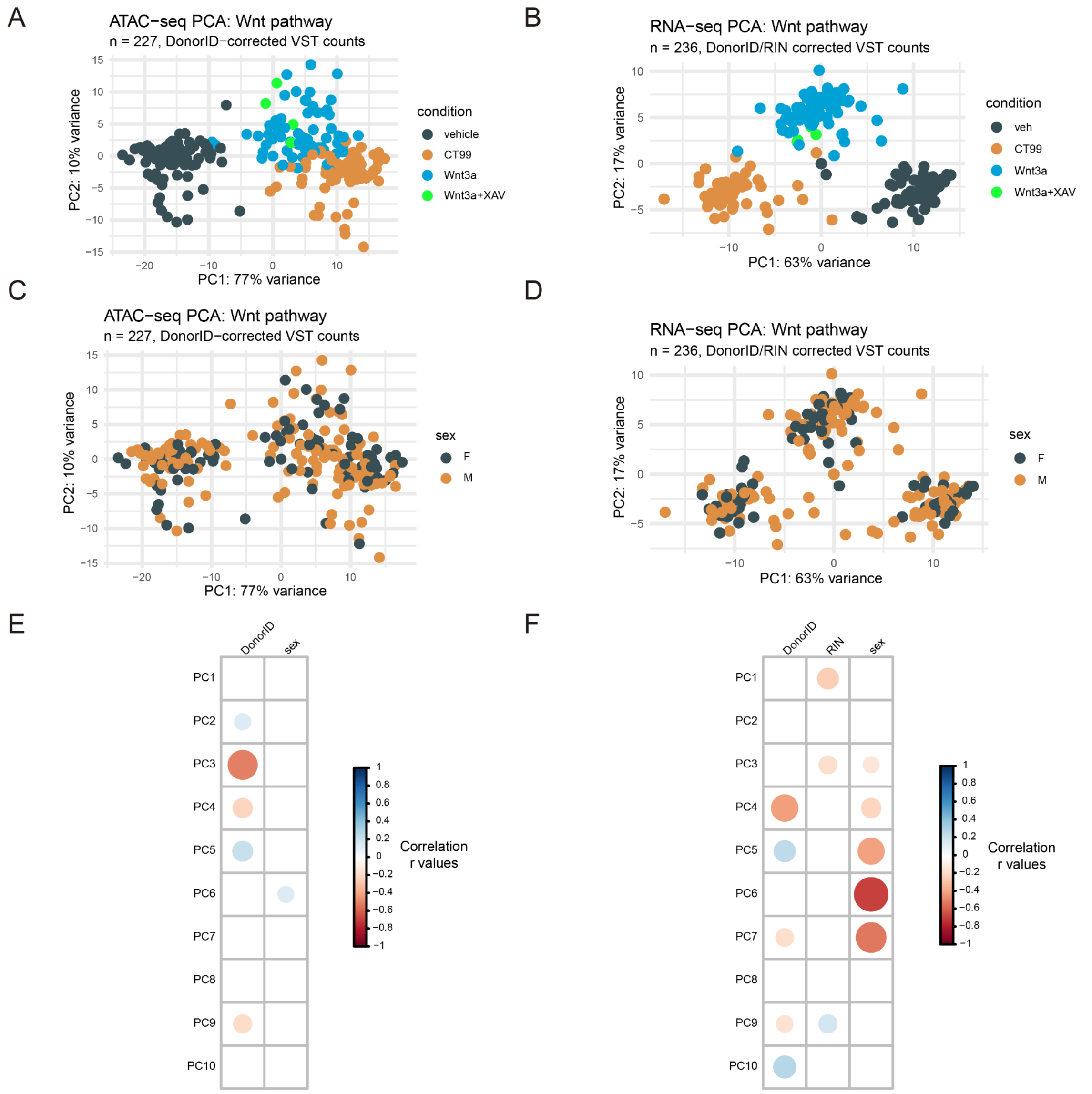

### Supplementary Figure 6| Number of differentially accessible regions and differentially expressed genes

Barplots showing the number of differentially accessible regions (WREs) (**A**) and differentially expressed genes (DEGs) (**B**). The reference indicates referenced condition (either vehicle or WNT3A). Sample size indicates paired samples between conditions. Percentage of increased (shrunken log2FC > 0.5 in dark blue, shrunken log2FC > 0 in blue) or decreased (shrunken log2FC < -0.5 in red, shrunken log2FC < 0 in pink) accessibility or expression are shown for each category. Nonsignificant peaks and genes (FDR-adjusted P >=0.1) are colored gray. Greater changes were observed in both chromatin accessibility and gene expression for CHIR, a potent Wnt activator, as compared to WNT3A, an endogenous Wnt ligand. Simultaneous WNT3A activation and inhibition via XAV as compared to WNT3A activation yielded few differentially expressed genes and chromatin elements, likely due to the low sample size. We also note that 93,853 peaks and 9,890 genes were filtered out (adjusted *P* value were set to *NA*) by DESeq2[2,3] as those have low mean read counts.

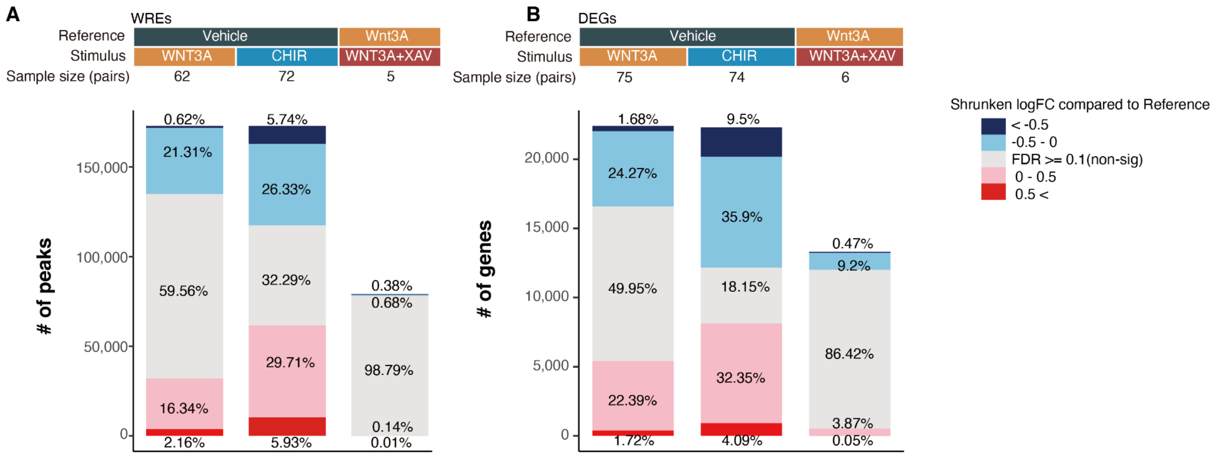

### Supplementary Figure 7| Wnt-pathway related transcription factor binding sites are enriched in upregulated WREs

The canonical Wnt-pathway downstream effectors, ß-catenin, LEF1, and TCF7L2 binding sites previously identified by ChIP-seq experiments in HEK cells[1,4] are more often overlapped with WREs opening due to WNT3A (**A**) or CHIR (**B**) than closing WREs. The figures show the percentage of overlaps with the indicated TF binding sites in either the 2,000 most upregulated peaks or the 2,000 most downregulated peaks based on shrunken logFC. Statistical significance was estimated using Fisher's exact test on the number of overlaps.

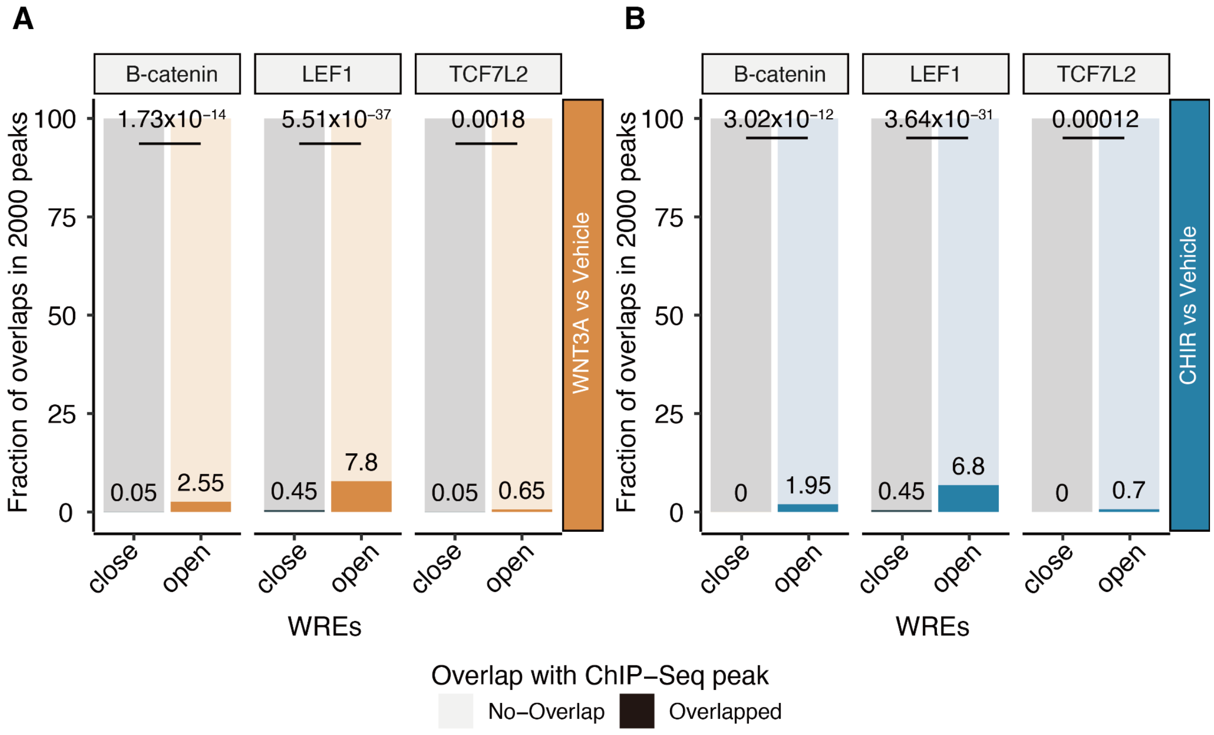

### Supplementary Figure 8| Stimulus-specific regulatory element-gene correlation

We identified stimulus-specific regulatory elements which regulate gene expression by correlating expression with chromatin accessibility. The number in the plot indicates significant and positively correlated peak-gene pairs. In total, 12,643 peak-gene pairs are detected only under the stimulus condition (highlighted in red), demonstrating new stimulus-specific regulatory elements.

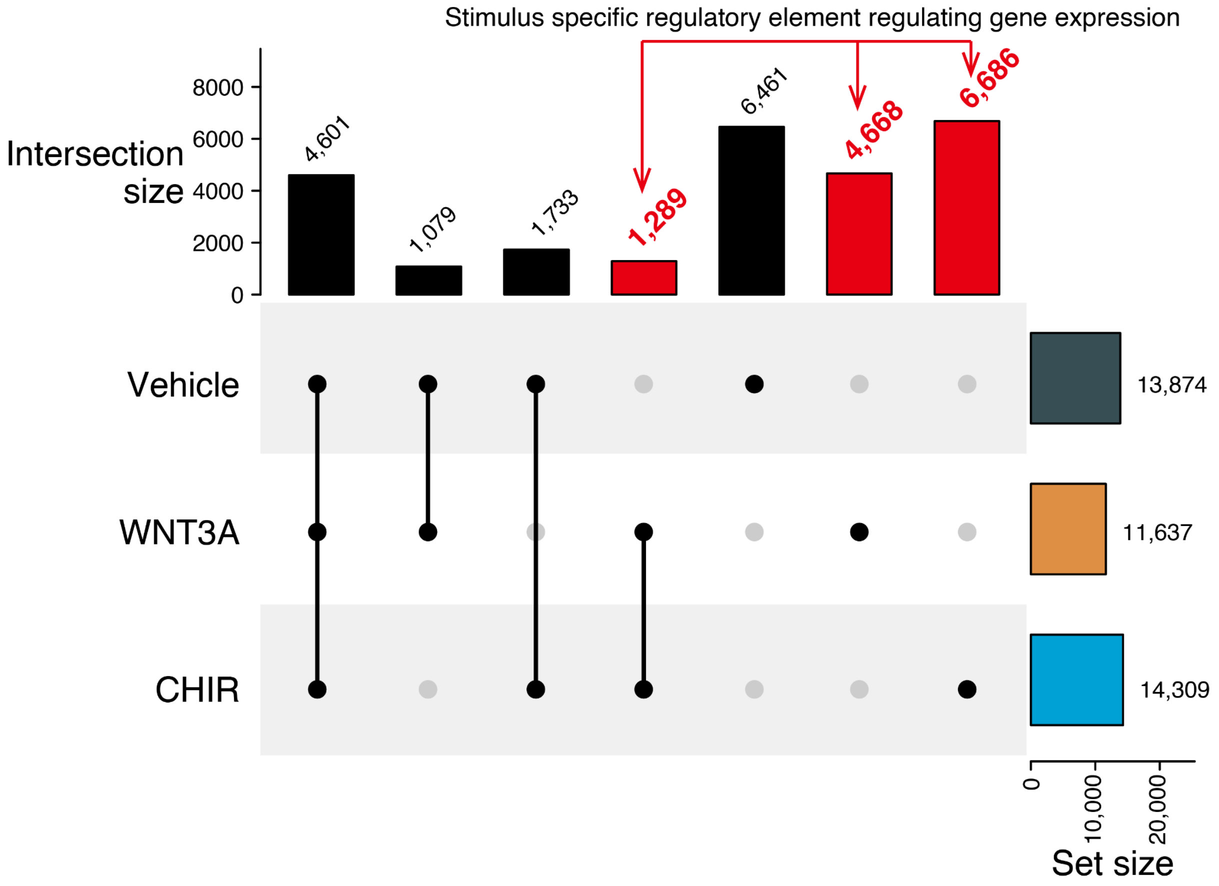

### Supplementary Figure 9| Opposing gene regulation by WNT pathway inhibition

(**A**) Volcano plot showing differential expressed genes between simultaneous excitation and inhibition of WNT3A (WNT3A+XAV) as compared to WNT3A stimulus alone. Note that LEF1, a downstream effector of the Wnt pathways, has decreased expression under inhibition of the Wnt pathway. (**B**) Among 1413 genes that are differentially expressed at BH-adj P < 0.1 in both comparisons (WNT3A+XAV vs WNT3A (6 pairs) and WNT3A vs Vehicle (75 pairs)), 70.8% (1001 genes), opposing changes in gene expression were observed. This result is consistent with most gene expression changes being caused by the stimulus of the Wnt pathway.

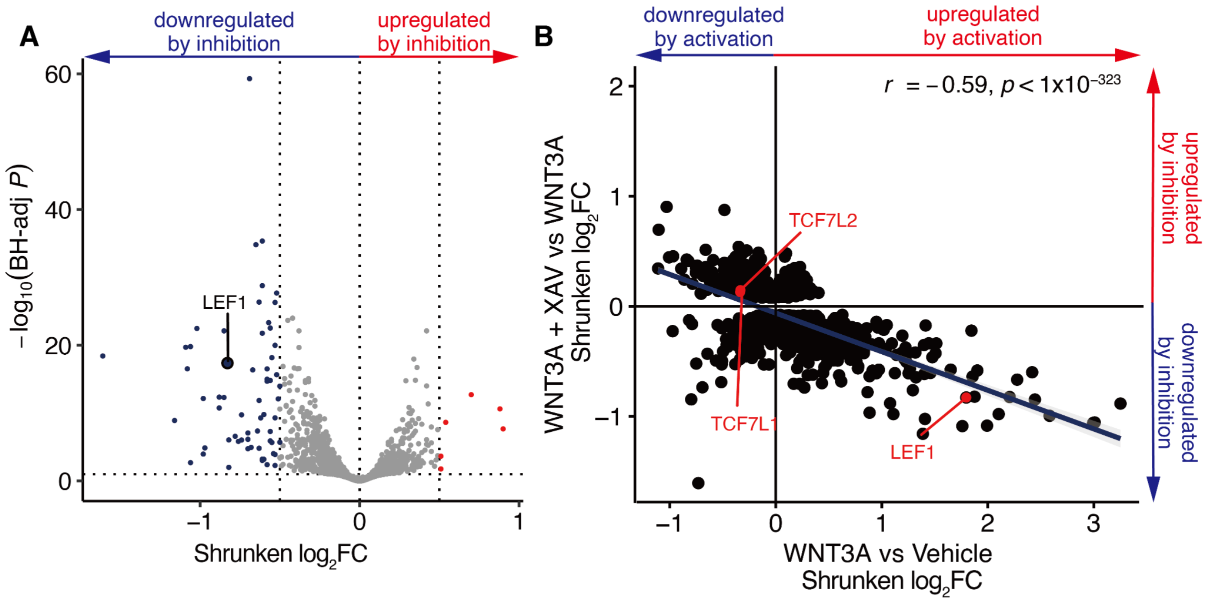

### Supplementary Figure 10| ca/eQTL effect sizes compared to previous data

We compared the effect size of ca-QTL (**A**) and eQTL (**B**) in vehicle condition to previous studies using the same hNPC cell lines with slightly larger sample size[5,6]. Strongly correlated effect sizes for unstimulated ca/eQTLs were found between the two datasets. In part **A**, the current study beta values are derived using RASQUAL as a scaled allelic ratio ranging from -0.5 to 0.5, which is not on the same scale as the beta values from Liang et al., which are calculated using a linear mixed effect model. One SNP showed opposite effect directionality across the caQTL datasets between the current and previous datasets (rs2076179). We suspect this was due to an error in allele assignment for that SNP in the previous analysis caused by sampling differences incorrectly changing the labeled minor allele for a SNP with allele frequency close to 0.50. We corrected this error for the one SNP prior to plotting.

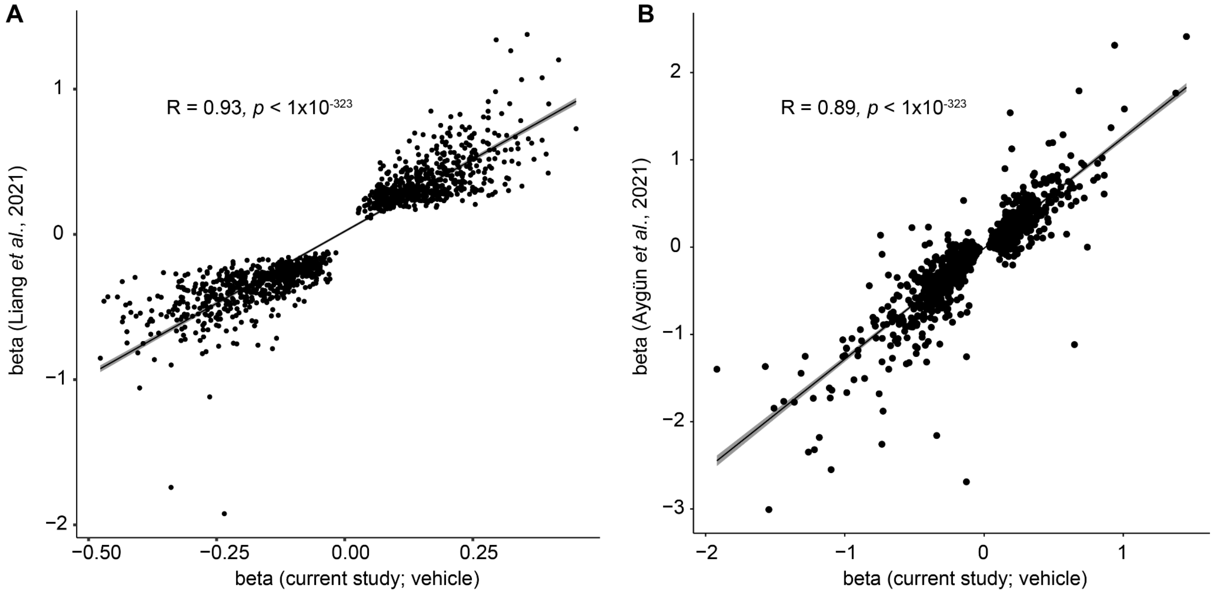

### Supplementary Figure 11| eQTL-caQTL overlaps

Overlaps between caQTLs and eQTLs were called where a caSNP was within 1Mb of an eGene and at least 1 significant SNP in both datasets were in LD $r^2 \geq 0.8$. The percentage of eQTLs (**A**) and caQTLs (**B**) overlapped within vehicle and WNT stimulus conditions are shown. The effect size of caQTL-eQTL sites which shared the same SNP position were compared within vehicle (**C**) and WNT stimulus conditions (WNT3A and CHIR shown in **D** and **E** respectively). SNPs selected to influence chromatin accessibility had relatively little overlap with those influencing gene expression likely because genetic variation affecting chromatin accessibility often does not lead to changes in gene expression (about <2% of caQTLs had a shared eQTL based on LD overlap ($r^2$ >= 0.8). However, SNPs selected to influence gene expression more often also influence chromatin accessibility in that ~27% of eQTLs have a shared caQTL within condition, a comparable number observed in our previous study (34.9% caQTL-eQTLs overlap[5]).

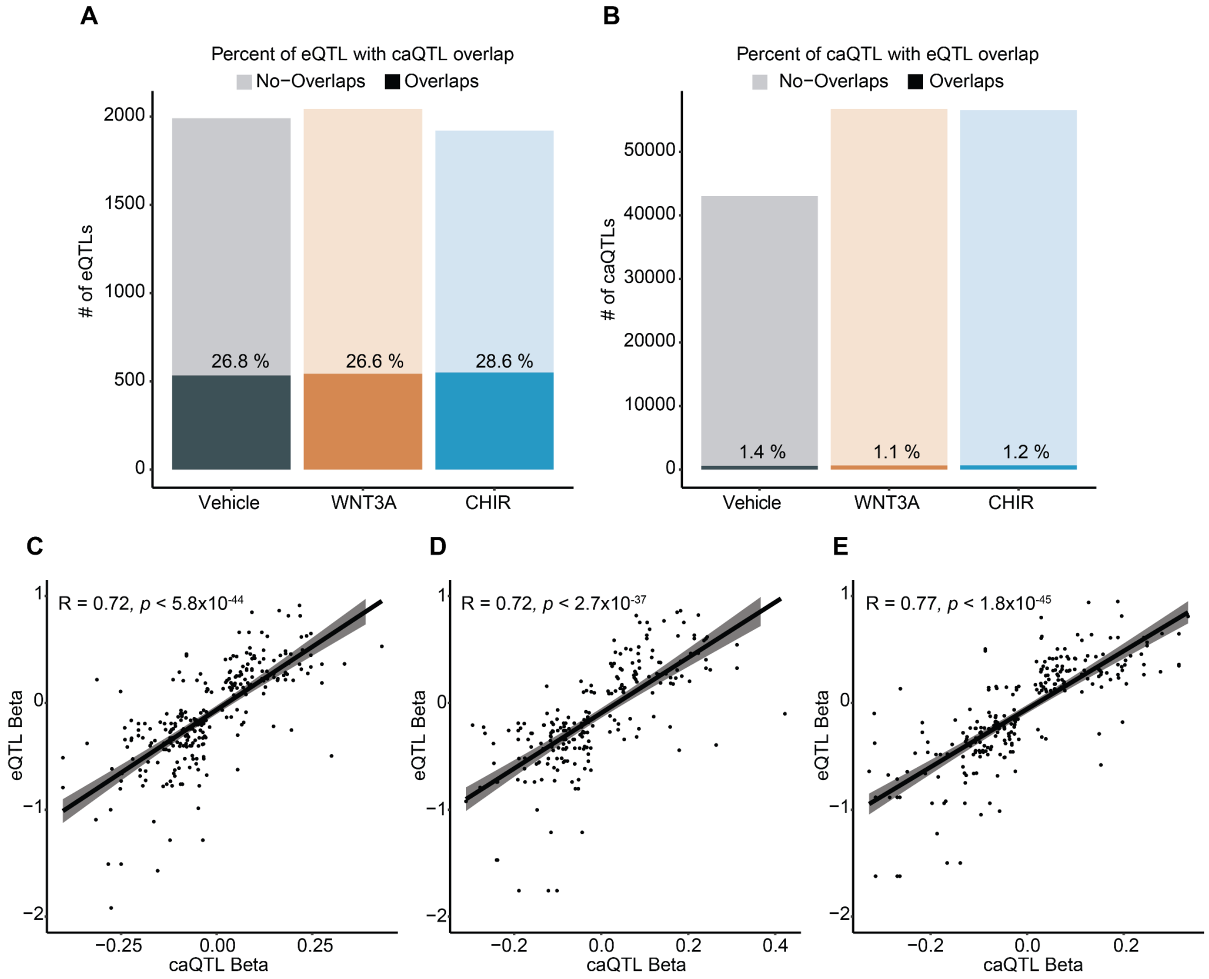

### Supplementary Figure 12| Effect size differences across conditions

We compared effect sizes of index caQTLs (**A**-**B**) / eQTLs (**C**-**D**) in stimulated conditions versus vehicle conditions (WNT3A or CHIR in (**A,C**) and (**B,D**), respectively). Dots in red indicate significant interaction effects were observed (r-QTLs). Error bars are standard errors of beta. We observed lower correlation in r-QTLs as compared to non-r-QTLs. We note that some SNPs are not tested for a baseline model due to low expression thus could not be plotted because no beta value was calculated.

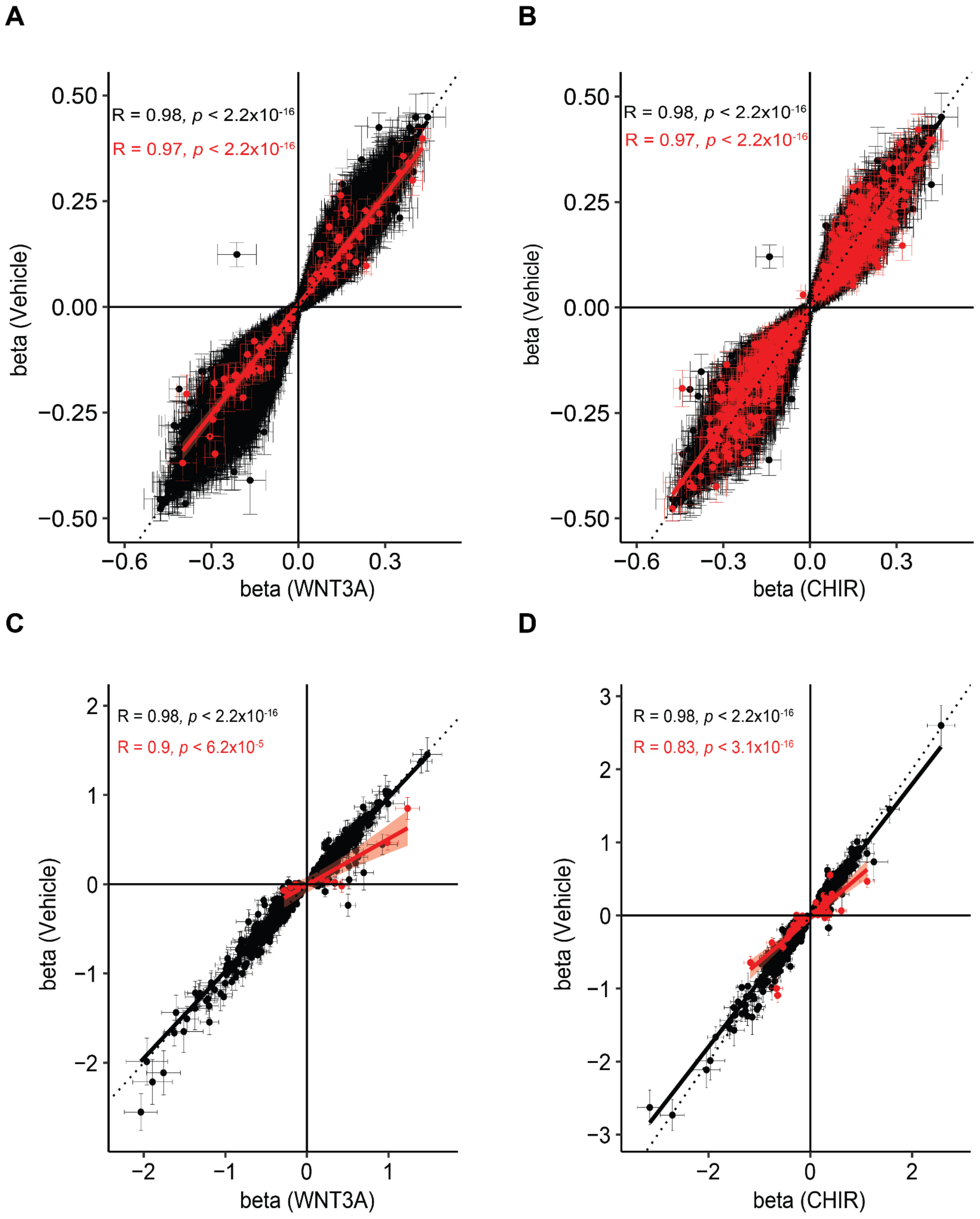

### Supplementary Figure 13| Response and Non−Response caPeaks enrichment within chromHMM Annotations

Enrichment of response and non-response caPeaks within chromHMM states defined within fetal brain male (E081)[7]. *, **, *** indicate enrichments of overlaps relative to the entire genome, evaluated with a binomial test, similar to the GREAT test[8], with *P*-values < 0.05, .01 and .001, respectively. We observed a significant enrichment of both response and non-response caPeaks within promoters, enhancers, and bivalent regions. TssA, TssFlnk, and Enh states showed a significant difference in overlap counts between response and non-response caPeaks (P = $3.9x10^{-5}$; $2.8x10^{-4}$; $7.4x10^{-7}$, respectively), as evaluated with Fisher's exact test. There was significantly less enrichment of response caPeaks in active TSS and enhancers as compared to non-response caPeaks, perhaps because these response caPeaks flag novel condition specific enhancers not annotated in post-mortem fetal brain tissue.

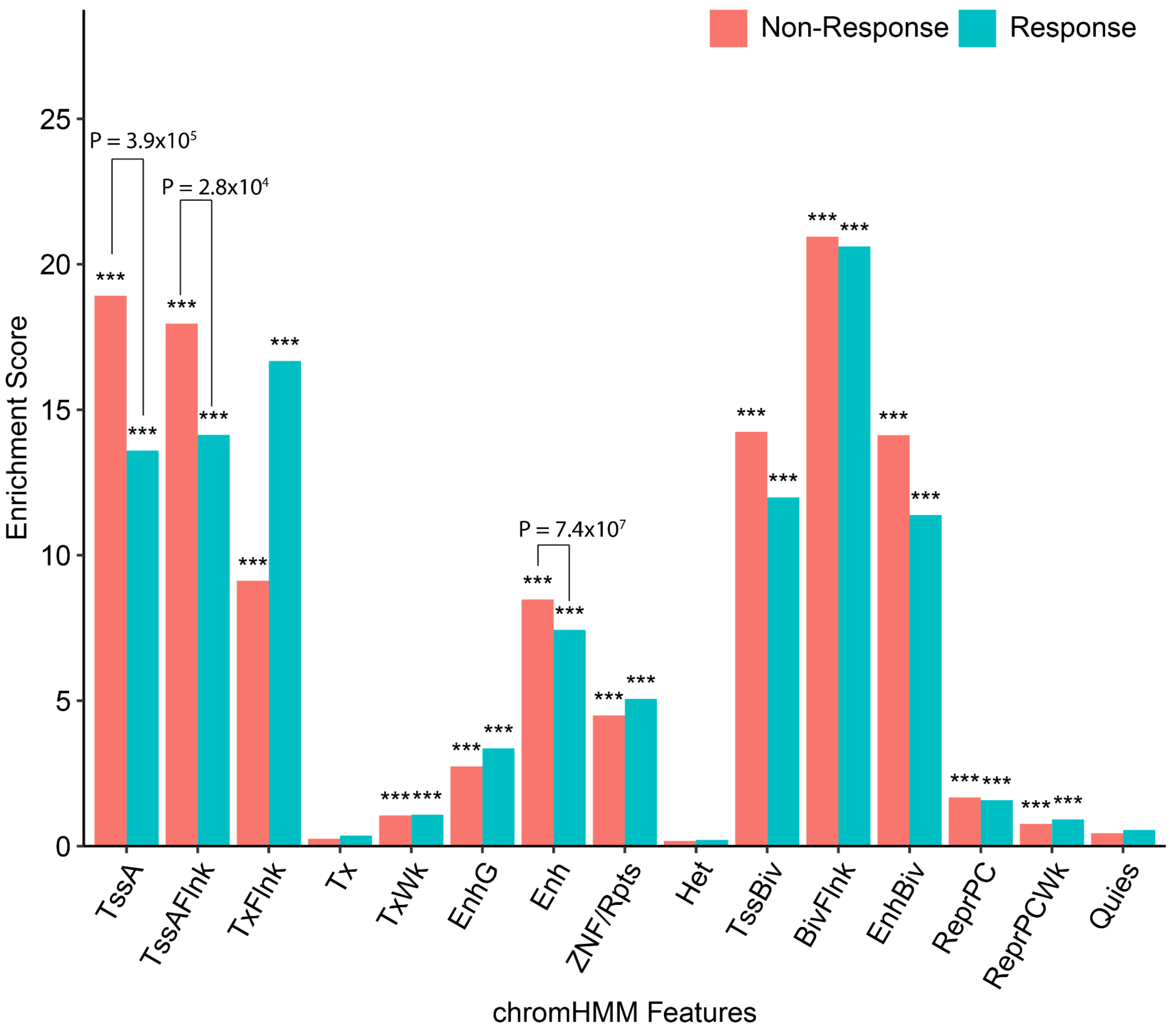

### Supplementary Figure 14| Context-specific caPeak overlaps HAQER near HAR1A/B

(**A**) Allelic effects of rs112675456 on chromatin accessibility (chr20:63101761-63102670), reveal a caQTL significant in CHIR stimulated condition but not vehicle. (**B**) Regional association plots at rs112675456, the index SNP for a caPeak-HAQER overlap near the transcription start sites of HAR1A/B. From top to bottom: Genomic coordinates, gene models, caQTL *P* values for vehicle and CHIR-stimulated conditions, ATAC-seq coverage showing differential chromatin accessibility, and HAQER.

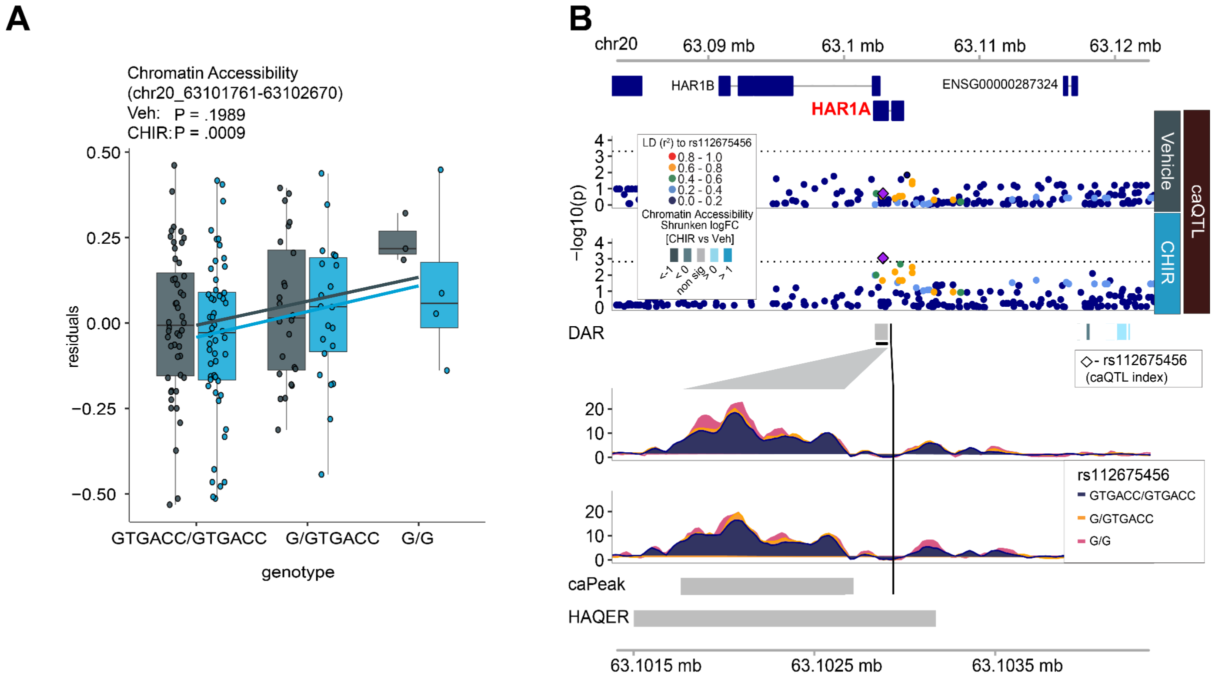

### Supplementary Figure 15| Colocalization of ANKRD44 r-eQTL, schizophrenia, and the volume of a hippocampal subfield (presubiculum body)

(**A**) Boxplot showing increased ANKRD44 gene expression by rs979020-T and the SNP a significant interaction effect. (**B**) P-P plot from schizophrenia GWAS vs eQTL colored by $r^2$ in this study population to rs979020, providing evidence for a colocalization. (**C**) Conditional analysis of schizophrenia GWAS was performed using GCTA-cojo tool with LD from the UKBB reference panel (White British). GWAS *P* value (upper panel) and post-conditional analysis *P* value (bottom panel) are shown. An absence of GWAS signal after conditioning on the r-eQTL index provides evidence for colocalization. (**D**, **E**) Similar to (**B**, **C**) but using GWAS for volume of a hippocampal subfield (presubiculum body; left hemisphere).

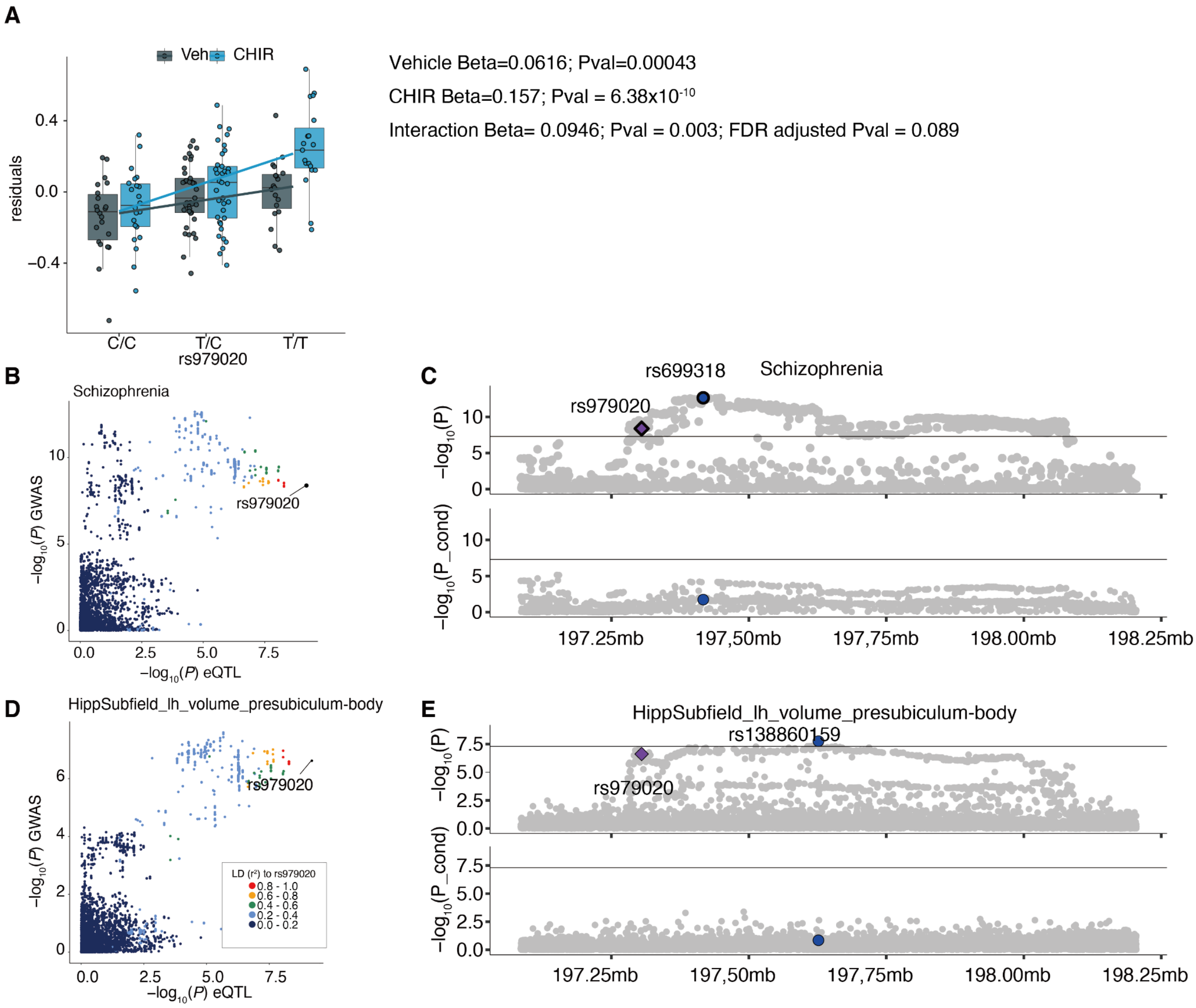

### Supplementary Figure 16| Colocalization of an r-caQTL in DPYSL5 region and average thickness of the isthmus Cingulate

(**A**) We detected an r-caQTL at rs1992311 (FDR-adjusted *P* value = 0.04). We present boxplots showing differential genetic effects on chromatin accessibility in stimulated versus vehicle conditions in a linked SNP within the peak that disrupts a motif (rs4665363-G). (**B**) P-P plot for isthmus Cingulate GWAS[9] vs caQTL colored by $r^2$ in our population to rs1992311, providing evidence for colocalization. (**C**) Boxplot showing decreased *DPYSL5* expression by rs4665363-G. (**D**) Conditional analysis on GWAS for average thickness of Isthmus Cingulate was performed using the UKBB reference panel (White British). GWAS *P* value (upper panel) and post-conditional analysis *P* value (bottom panel) are shown. GCTA-cojo identified collinearity between rs4665363 and rs3820823 ($r^2$>0.9) thus both SNPs are not shown in the bottom panel. A decrease in GWAS signal after conditioning on the r-caQTL provides evidence for colocalization. (**E**) Logo plot predicting disruption of the Pou5f1::Sox2 motif by rs4665363-G. (**F**) Coverage plots of the peak showing the location of rs4665363 and a TCF/LEF motif present in the peak.

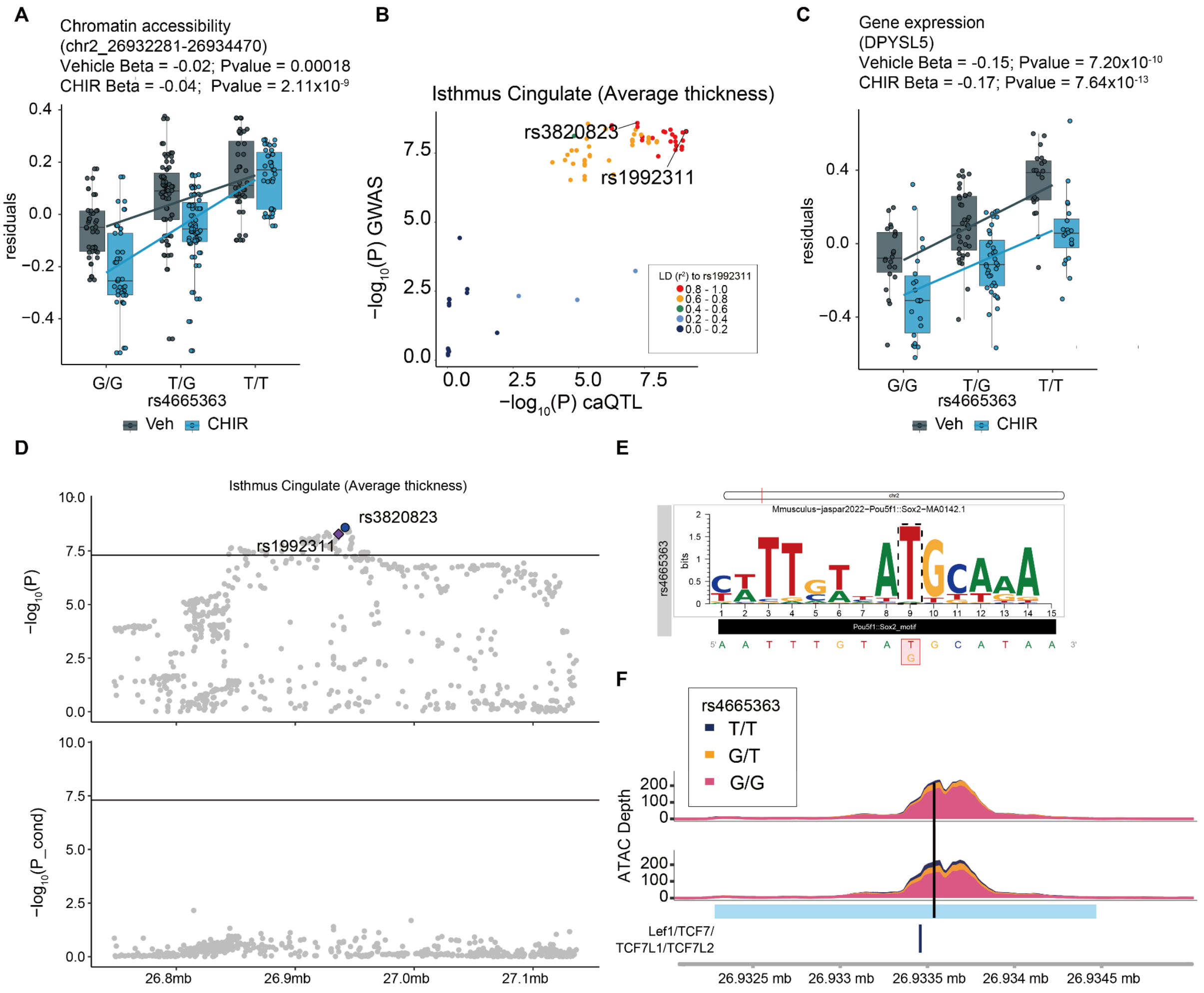

### Supplementary Figure 17| Shared and context-specific QTL GWAS overlaps confirmed by eCAVIAR

Colocalization of eQTLs/caQTLs and brain-related GWAS phenotypes tested by eCAVIAR. (**A**) A cumulative number of colocalized eGenes/caPeaks after colocalization analysis by eCAVIAR and (**B**) shared/distinct colocalization in each condition. The use of stimulated conditions increased the number of brain-trait associated genes by 75.6% and peaks by 77.0%.

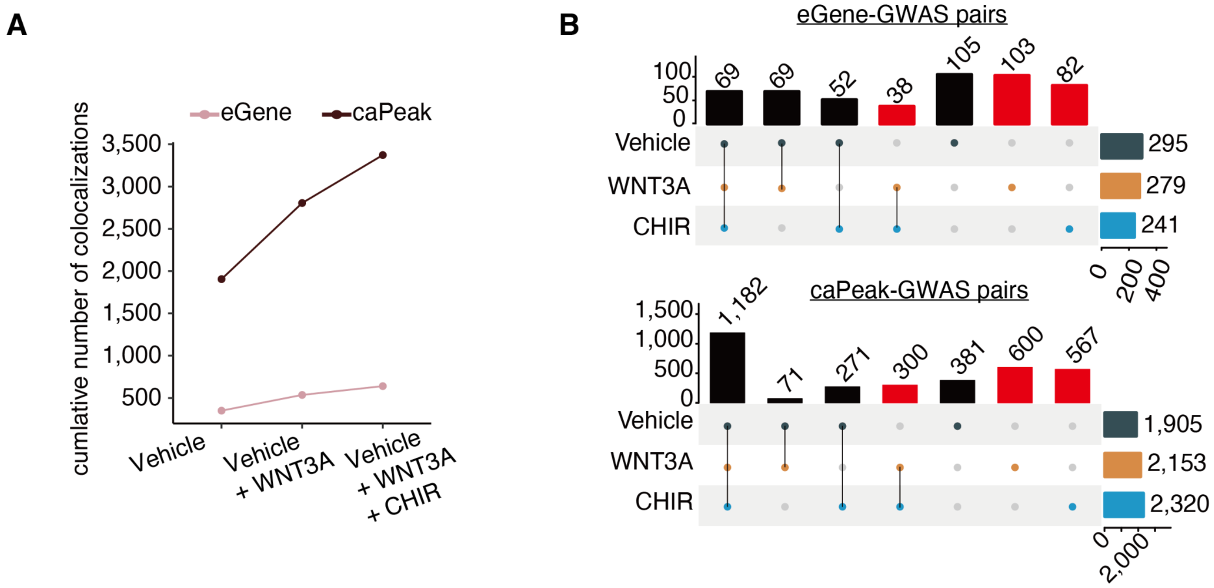

### Supplementary Figure 18| Stimulus-specific GWAS colocalization of *FADS3* and Bipolar disorder

(**A**) GWAS and QTL locus plots in the region of *FADS3*. GWAS index SNP rs174592-A (protective allele) of bipolar disorder (BP) is in high LD with index caQTL (rs99780, $r^2$ = 0.86 in EUR, 0.85 in this study population) for RE (chr11:61850491-61851800) and *FAD3* index eQTL (rs174583, $r^2$ = 0.89 in EUR, $r^2$ = 0.78 in this study population). Rs174592-A is located within the peak and predicted to disrupt a STAT1::STAT2 transcription factor binding site motif. (**B**) Rs174592-A decreases chromatin accessibility of this WRE (Beta = -0.10; P = 6.81 x $10^{-13}$, left), and expression of *FADS3* (Beta = -0.13 P = 2.95 x $10^{-7}$, right) in CHIR condition. (**C**) Differential chromatin accessibility and *FADS3* gene expression between CHIR and Vehicle. (**D**) VST normalized expression counts of STAT1 and STAT2 are shown. Shrunken LFC and FDR-adjusted P values were estimated for 78 pairs. In the boxplots, chromatin accessibility or gene expression are colored by condition (gray: vehicle, blue:CHIR). (**E**) Schematic of FADS3 regulation through STAT1::STAT2 binding. Our data suggests characterization of the colocalized putative BP risk SNP (rs174592) as a functional variant regulating *FADS3* expression through differences in STAT1::STAT2 binding, which is inhibited during Wnt stimulation.

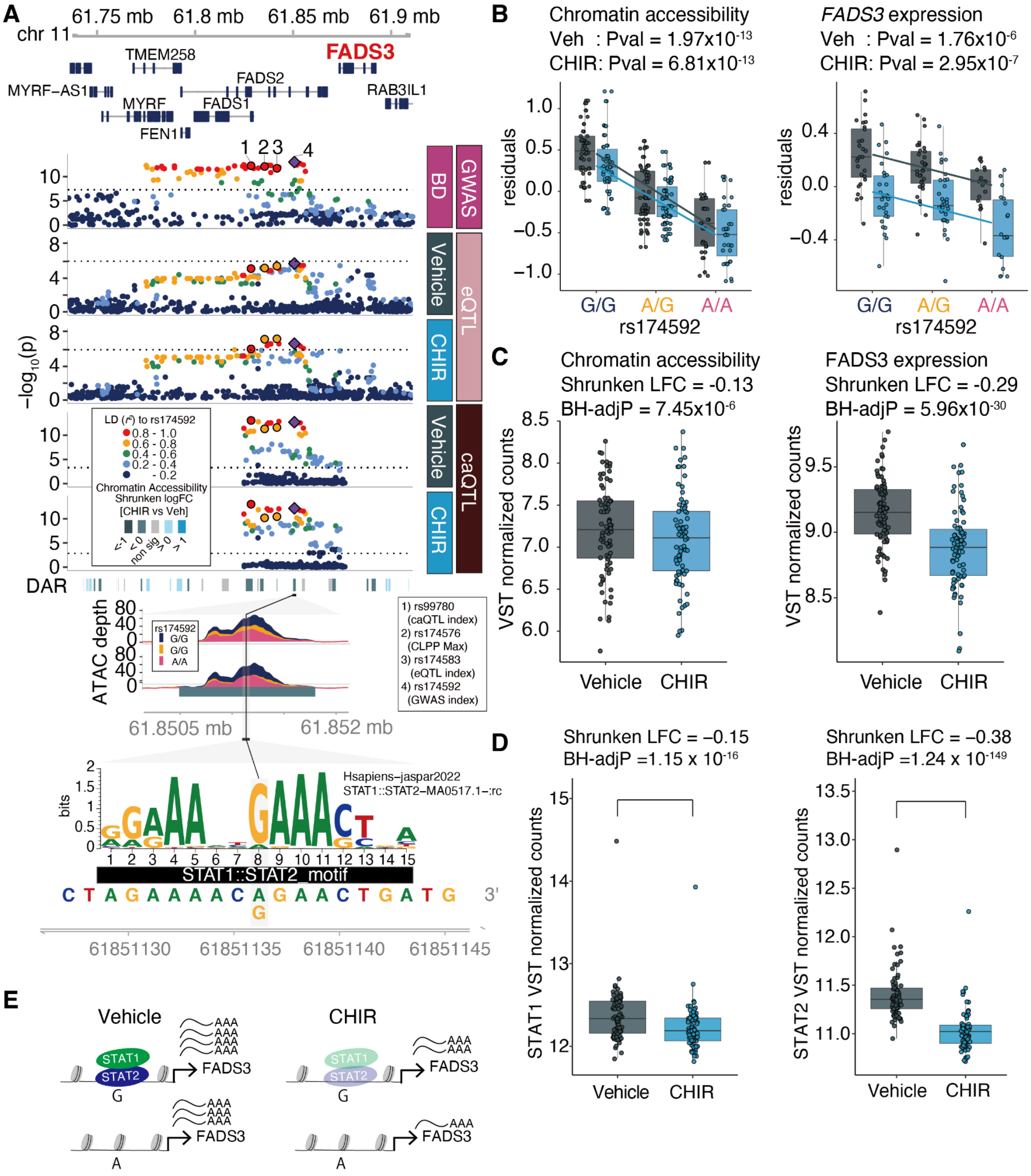

### Supplementary Figure 19| Stimulus-specific *ENO4* eQTL colocalizing with regional cortical surface area GWAS

(**A**) Chromatin accessibility (chr10:116973711-116975130; left) and *ENO4* gene expression regulated by rs11197861. eQTL of *ENO4* was detected only in the stimulated condition (**A**, **B**). The eQTL is colocalized with several brain-related traits including surface area of Insula (**B**, **C**). (**B**) *P* values from Insula surface area GWAS, eQTL, caQTL for vehicle, CHIR condition are shown respectively. ETV2:FOXI1 motif is disrupted by the rs11197861-T located in the peak, which may result in decreased chromatin accessibility and *ENO4* gene expression. (**C**) Z scores from regional cortical surface area[9]. In addition to Insula, surface area of precentral and precuneus are also associated with this SNP (indicated by asterisks; $p < 5 \times 10^{-8}$).

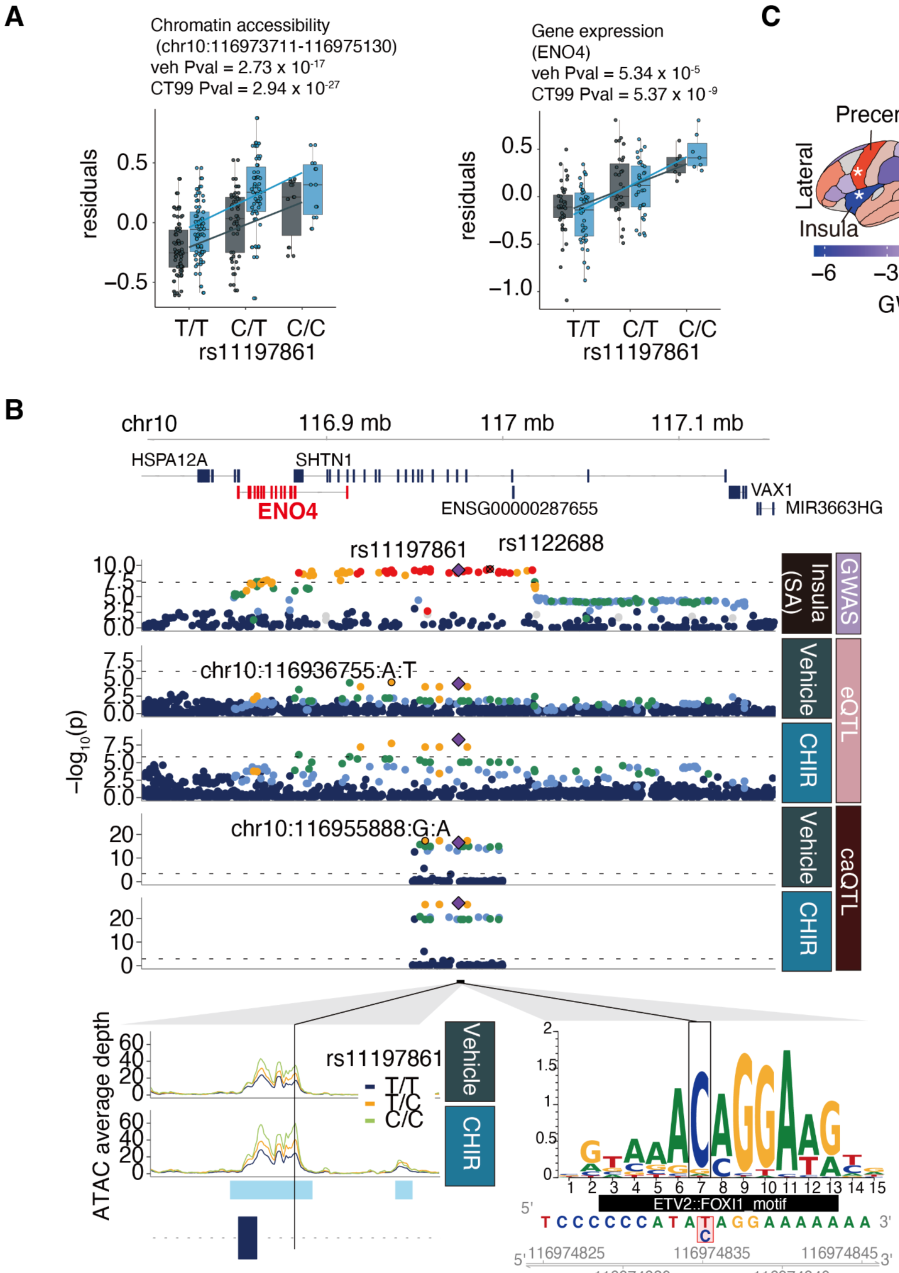

### Supplementary Figure 20| Optimizing control for known and unknown technical confounding in eQTL

We tested different numbers of PC variables to determine the number of PCs to include for correcting expression values prior to running limix_qtl. (**A**) The plot shows the number of discovered eGenes with respect to the number of PCs at FDR-adjusted P < 0.1. and the number of overlapping found eGenes across models (**B-D**). To identify independent eQTLs, we repeated eQTL mapping by including the index eSNP in the association model until no SNP passed the genome-wide threshold (FDR-adjusted P < 0.1). In table (**E**), we show the number of SNPs discovered in each round of conditional associations.

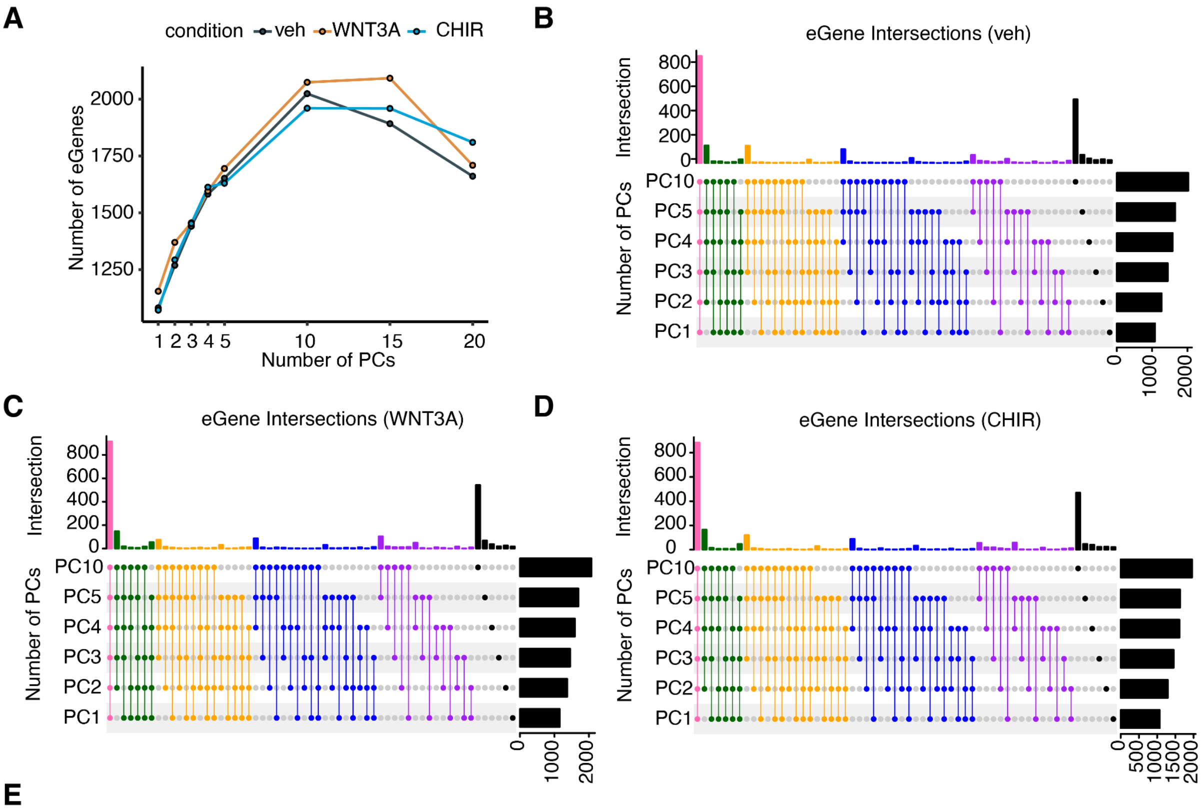


**E**

| # of eSNPs per Gene | | |
|---|---|---|
| **Round** | **# eGenes** | **condition** |
| 1 | 2024 | Vehicle |
| 2 | 65 | Vehicle |
| 1 | 2074 | Wnt3A |
| 2 | 72 | Wnt3A |
| 3 | 1 | Wnt3A |
| 1 | 1960 | CHIR |
| 2 | 65 | CHIR |
| 3 | 1 | CHIR |

### Supplementary Tables

S1 - S4, S6-S19 are provided in Excel files.

#### Supplementary Table 1| Differentially accessible peaks

A. WNT3A vs Vehicle; B. CHIR vs Vehicle; C. WNT3A+XAV vs WNT3A

| Column Name | Description |
|---|---|
| Chromosome | chromosome |
| start | peak start position (hg38) |
| end | peak start position (hg38) |
| baseMean | average of the normalized count values |
| log2FoldChange | shrunken log2Fold Change |
| lfsSE | standard error of Shrunken log2Fold Change |
| pvalue | p value |
| padj | BH-adjusted Pvalue |

#### Supplementary Table 2| TF motif enrichment analysis

A. WNT3A vs Vehicle; B. CHIR vs Vehicle

| Column Name | Description |
|---|---|
| TFname | TF name |
| TFID | TF ID |
| estimate | coefficient from logistic regression |
| estimate_err | coefficient standard error from logistic regression |
| z | z score = estimate/estimate_err |
| pval | pvalue from logistic regression |
| padj | BH-adjsted Pvalue |

#### Supplementary Table 3| Differentially expressed genes

A. WNT3A vs Vehicle; B. CHIR vs Vehicle; C. WNT3A+XAV vs WNT3A

| Column Name | Description |
|---|---|

| | |
|---|---|
| gene_id | Ensembl ID |
| gene_name | HGNC gene name |
| gene_type | protein_coding/lncRNA |
| chromosome | chromosome |
| start | collapsed gene start position |
| end | collapsed gene end position |
| strand | strand |
| baseMean | average of the normalized count values |
| log2FoldChange | shrunken log2Fold Change |
| lfcSE | standard error of Shrunken log2Fold Change |
| pvalue | p value |
| padj | BH-adjusted Pvalue |

### Supplementary Table 4| Pathway enrichment analysis

A. WNT3A vs Vehicle; B. CHIR vs Vehicle
Details can be found on the gprofiler website (https://biit.cs.ut.ee/gprofiler/page/apis)

| **Column Name** | **Description** |
|---|---|
| query | DEG direction (up-regulated/down-regulated/all) |
| source | KEGG / REAC(REACTOME) |
| term_id | pathway ID |
| term_name | pathway name |
| p_value | Hypergeometric p-value after correction for multiple testing (BH) |
| query_size | number of genes in the query |
| intersection_size | number of genes in the query annotated to the corresponding term |
| term_size | number of genes annotated to the term |
| effective_domain_size | total number of genes in the background that is used for Hypergeometric test |
| GeneRatio | interaction_size/query_size |

#### Supplementary Table 5| Summary of peak-gene correlation

| Condition | Pearson's Correlation Coefficient | | | % of positively correlated pairs |
|---|---|---|---|---|
| | max r | mean(abs.r) | min r | |
| Vehicle | 0.95 | 0.50 | -0.78 | 80.4% |
| WNT3A | 0.95 | 0.53 | -0.83 | 83.4% |
| CHIR | 0.97 | 0.50 | -0.75 | 83.6% |

#### Supplementary Table 6| Peak-gene pairs

A. WNT3A vs Vehicle; B. CHIR vs Vehicle

| Column Name | Description |
|---|---|
| chromosome | chromosome position of peak |
| start | start position of peak |
| end | end position of peak |
| gene_id | Ensembl ID |
| gene_name | HGNC gene name |
| distance | distance from TSS to center of peak |
| coefficient | coefficient in a linear model |
| fdr | BH-adjusted P value |
| (interaction coefficient) | Interaction coefficient in a linear model (WNT3A/CHIR vs Vehicle) |
| (fdr_interact) | BH-adjusted P value for interaction test (WNT3A/CHIR vs Vehicle) |

#### Supplementary Table 7| Disease enrichment analysis results(DEG)

Details can be found in DisGeNet (https://www.disgenet.org/home/).

| Column Name | Description |
|---|---|

| | |
|---|---|
| status | DEG category (all/up/down/nonsig) |
| cond | condition (WNT3A/CHIR/nonsig) |
| ID | phenotype ID |
| Description | phenotype description |
| source | “CURATED” |
| BgRatio | number of genes associated to the disease and number of background gene in database |
| pvalue | Fisher's exact test P value |
| FDR | BH-adjusted P value |
| disease_class | MESH tree branch class |
| disease_class_name | MESH tree branch name |
| disease_semantic_type | UMLS® semantic types |
| shared_symbol | shared gene symbol associated to the disease |

### Supplementary Table 8| Summary of GWASs used in this study

| **Column Name** | **Description** |
|---|---|
| GWAS Name | GWAS phenotype |
| Ref | Reference |

### Supplementary Table 9| Partitioned heritability analysis results

| **Column Name** | **Description** |
|---|---|
| Annotation type | Tissue / Cell / WNT stimulation |
| Data set | WREs used for LDSC |
| GWAS phenotype | GWAS phenotype |
| Coefficient | coefficient from LDSC model |
| Coefficient stderr | standard error of Coefficient |

| | |
|---|---|
| Coefficient P | One-sided Pvalue from Coefficient Z score |
| BH-adjusted P | BH-adjusted Pvalue |

#### Supplementary Table 10| List of caQTL - caPeak

| **Column Name** | **Description** |
|---|---|
| Feature | peak ID |
| rs.ID | SNP ID |
| Chromosome | chromosome position of peak/snp |
| SNP.Position | SNP position |
| Ref.Allele | reference allele |
| Alt.Alele | alternative allele (the effect allele) |
| Allele.Frequency | allele frequency of reference allele |
| HWE.Chi_Square.Statistic | Hardy-Weinberg chi-square statistic |
| Imputation.Quality..IA. | imputation quality score |
| Log_10.Benjamini.Hochberg.Q.value | $\log_{10}$Benjamini-Hochberg q-value |
| Chi.square.statistic..2.x.log.Likelihood.ratio. | chi square statistic (2 x log-likelihood ratio) |
| Effect.Size | cis-regulatory effect parameter (pi) |
| Sequencing.mapping.error.rate..Delta. | sequencing/mapping error rate (Delta) |
| Reference.allele.mapping.bias..Phi. | reference allele mapping bias (Phi) |
| Overdispersion | overdispersion estimate |
| SNP.ID.within.the.region | ID of snp within tested region |
| No..of.feature.SNPs | total number of snps tested within the feature |
| No..of.tested.SNPs | total number of snps tested within cis window |
| No..of.iterations.for.null.hypothesis | number of iterations for null hypothesis |
| No..of.iterations.for.alternative. | number of iterations for alternative hypothesis |

| | |
|---|---|
| hypothesis | |
| Random.location.of.ties | location of snp used as lead if tie occurs |
| Log.likelihood.of.the.null.hypothesis | log-likelihood of the null hypothesis |
| Convergence.status | job outputs status (0=success) |
| Squared.correlation.between.prior.and.posterior.genotypes..fSNPs. | squared correlation between prior and posterior genotypes (fSNPs; feature snps) |
| Squared.correlation.between.prior.and.posterior.genotypes..rSNP. | squared correlation between prior and posterior genotypes (rSNP; single cis-regulatory snp) |
| PValue | p value |
| Distance_from_PeakCenter | distance of tested snp from the center of the peak |

#### Supplementary Table 11| List of eQTL - eGene

| **Column Name** | **Description** |
|---|---|
| gene_id | Ensembl ID |
| gene_chr | chromosome position of gene |
| gene_start | start position of gene |
| gene_end | end position of gene |
| gene_name | HGNC gene name |
| feature_strand | strand (+/-) |
| snp_id | snp ID |
| snp_chr | chromosome position of SNP |
| snp_pos | SNP position |
| Assessed Allele | Assessed (also called effect) allele |
| maf | minor allele frequency |
| beta | beta estimate of main effect |
| beta_se | standard error of beta estimate |
| empirical_feature_p_value | permutation p value |

| | |
|---|---|
| p_value | p value |
| global_adj_p_value | BH-adjusted P value |
| iteration | iteration (1 = primary) |

### Supplementary Table 12| caSNP TF motif disruption (in peak)

| Column Name | Description |
|---|---|
| Chr | chromosome position |
| snpPos | snp position |
| strand | strand (+/-) |
| SNP_id | SNP ID |
| REF | reference allele |
| ALT | alternative allele |
| geneSymbol | motif gene symbol |
| dataSource | motif data source |
| providerId | JASPAR matrix ID |
| effect | disruption effect strength |
| motifPosition_start | start position of disrupted motif |
| motifPosition_end | end position of disrupted motif |

### Supplementary Table 13| List of response-caQTLs

| Column Name | Description |
|---|---|
| Feature | peak ID |
| SNP | snp ID |
| PValue | p value |
| Interaction_Beta | interaction effect |

| | |
|---|---|
| Chr | chromosome position |
| Position | SNP position |
| global | BH-adjusted Pvalue |
| Peak_SNP | combined peak ID and snp ID |
| Ref | reference allele |
| Alt | alternative allele |
| RASQUAL_PValue_Veh | p value from vehicle RASQUAL output |
| RASQUAL_PValue_Condition | p value from condition RASQUAL output |
| Pi_Veh | cis-regulatory effect parameter (pi) from vehicle RASQUAL output |
| EffectSize_Veh | beta estimate of snp effect in vehicle |
| Pi_Condition | cis-regulatory effect parameter (pi) from condition RASQUAL output |
| EffectSize_Condition | beta estimate of snp effect in condition |

### Supplementary Table 14| List of response-eQTLs

A. WNT3A vs Vehicle, B. CHIR vs Vehicle

| **Column Name** | **Description** |
|---|---|
| gene_name | HGNC gene name |
| chromosome | chromosome position |
| start | start position of gene |
| end | end position of gene |
| snp_id | SNP ID |
| Assessed Allele | assessed allele |
| maf | minor allele (assessed allele) frequency |
| beta | interaction effect |
| beta_se | standard error of interaction effect |
| p_value | P value of interaction effect |

| | |
|---|---|
| padj | BH-adjusted Pvalue |

#### Supplementary Table 15| TFBS motif enrichment in response-caPeaks

A. WNT3A vs Vehicle response-caPeaks, B. CHIR vs Vehicle response-caPeaks

| **Column Name** | **Description** |
|---|---|
| TF_name | Name of transcription factor binding site motif (JASPAR 2022) |
| TFID | JASPAR 2022 ID for transcription factor binding site |
| pval | P-value for logistic regression fit from r/glm() indicates significance of enrichment in response vs. non-response caPeaks |
| estimate | Estimated coefficient for effect of caPeak type (response vs. non-response) on TFBS motif presence in the caPeak |
| estimate_error | Standard error of estimates |
| z | Scaled estimates (estimate/estimate_error) |
| fdr | BH-adjusted Pvalue |

#### Supplementary Table 16| caQTL Regional Overlaps

| **Column Name** | **Description** |
|---|---|
| CHROM | chromosome position |
| START | start position of HAQER |
| END | End position of HAQER |
| NAME | HAQER ID |
| Feature | caPeak ID |
| Haq_Index | Index of HAQER |
| Wnt_Index | Index of caPeak |
| Region | Overlap region type (HAQERs/HARs) |
| Response_caPeak | * indicates response caPeak |

#### Supplementary Table 17| caQTL-eQTL colocalizations

| Column Name | Description |
|---|---|
| gene_chr | chromosome position (eQTL) |
| gene_start | start position of gene |
| gene_end | end position of gene |
| gene_name | HGNC gene name |
| snp_id | SNP ID (eQTL) |
| Assessed Allele | assessed allele (eQTL) |
| maf | minor allele (assessed allele) frequency (eQTL) |
| beta | interaction effect (eQTL) |
| beta_se | standard error of interaction effect (eQTL) |
| p_value | P value of interaction effect (eQTL) |
| global_adj_p_value | BH-adjusted Pvalue (eQTL) |
| Feature | peak ID |
| rs.ID | SNP ID (caQTL) |
| Chromosome | chromosome position (caQTL) |
| SNP.Position | SNP position (caQTL) |
| Ref.Allele | reference allele (caQTL) |
| Alt.allele | alternative allele (caQTL) |
| Effect.Size | cis-regulatory effect parameter (pi) |
| PValue | p value (caQTL) |
| Distance_from_PeakCenter | distance of tested SNP from the center of the peak |
| R2 | *r2* between eQTL and caQTL SNP |

#### Supplementary Table 18| GWAS colocalization results (caQTL)

| Column Name | Description |
|---|---|
| chr | chromosome |
| Peak | peak ID |
| Wnt_Pos | caQTL index position |
| Wnt_SNP | caQTL SNP ID |
| GWAS_Pos | GWAS index Position |
| GWAS_SNP | GWAS SNP ID |
| R2_CurrentStudy | $r^2$ between GWAS index SNP and eQTL index SNP estimated in this study population |
| R2_1kgEUR | $r^2$ between GWAS index SNP and eQTL index SNP estimated in 1KG EUR population |
| GWASID | GWAS phenotype(s) |

### Supplementary Table 19| GWAS colocalization results (eQTL)

| Column Name | Description |
|---|---|
| condition | stimulus condition (veh/WNT3A/CHIR) |
| gene | HGNC gene name |
| chr | chromosome |
| WntPos | eQTL index position |
| GWAS Pos | GWAS index Position |
| r2_current_study | $r^2$ between GWAS index SNP and eQTL index SNP estimated in this study population |
| r2_1kgEUR | $r^2$ between GWAS index SNP and eQTL index SNP estimated in 1KG EUR population |
| WntindexSNP | eQTL index SNP id |
| Pheno2indexrsID | Overlapping GWAS index SNP rsid |
| gtxSigInLocus_Brain_Cortex | TRUE: significant GTEx bulk tissue eQTL detected within $r^2 > 0.6$ of eQTL index SNP<br>FALSE: significant GTEx bulk tissue eQTL NOT detected within $r^2 > 0.6$ of eQTL index SNP |
| gtxSigInLocus_Brain_Frontal_ | As above for GTEx Brain_Frontal_Cortex_BA9 tissue |

| | |
|---|---|
| Cortex_BA9 | |
| gtxSigInLocus_Adipose_Subcutaneous | As above for Adipose_Subcutaneous |

### Supplemental Information References
